# Entrained audiovisual speech integration implemented by two independent computational mechanisms: Redundancy in left posterior superior temporal gyrus and Synergy in left motor cortex

**DOI:** 10.1101/202911

**Authors:** Hyojin Park, Robin A. A. Ince, Philippe G. Schyns, Gregor Thut, Joachim Gross

**Affiliations:** Institute of Neuroscience and Psychology, University of Glasgow, Glasgow, United Kingdom; Institute for Biomagnetism and Biosignalanalysis, University of Munster, Munster, Germany

## Abstract

Information integration is fundamental to many aspects of human behavior, and yet its neural mechanism remains to be understood. For example, during face-to-face communication we know that the brain integrates the auditory and visual inputs but we do not yet understand where and how such integration mechanisms support speech comprehension. Here we show that two independent mechanisms forge audiovisual representations for speech comprehension in different brain regions. With a novel information theoretic measure, we found that theta (3-7 Hz) oscillations in the posterior superior temporal gyrus/sulcus (pSTG/S) code speech information that is common (i.e. redundant) to the auditory and visual inputs whereas the same oscillations in left motor and inferior temporal cortex code synergistic information between the same inputs. Importantly, redundant coding in the left pSTG/S and synergistic coding in the left motor cortex predict behavior - i.e. speech comprehension performance. Our findings therefore demonstrate that processes classically described as *integration* effectively reflect independent mechanisms that occur in different brain regions to support audiovisual speech comprehension.

## Introduction

While engaged in a conversation, we effortlessly integrate auditory and visual speech information into a unified perception. Such integration of multisensory information is a key aspect of audiovisual speech processing that has been extensively studied [1–4]. Studies of multisensory integration have demonstrated that, in face-to-face conversation, especially in adverse conditions, observing lip movements of the speaker can improve speech comprehension [4–6]. In fact, lip movements during speech typically contain sufficient information to understand speech without corresponding auditory information [1, 7].

Turning to the brain, we know that specific brain regions are involved in audiovisual integration. Especially the superior temporal gyrus/sulcus (STG/S) responds to conjunction of auditory and visual stimuli and its disruption leads to reduced McGurk fusion [8–12]. However, these classic studies present two shortcomings. First, their experimental designs typically contrasted two conditions: unisensory (i.e. audio or visual cues) and multisensory (congruent or incongruent audio and visual cues). However, such contrast does not dissociate effects of integration *per se* from those arising from differences in stimulation complexity (i.e. one or two sources) – e.g. possible modulations of attention, cognitive load and even arousal. A second shortcoming is that previous studies typically investigated (changes of) regional activation and not information integration between audiovisual stimuli and brain signals. Here, we address these two shortcomings and study the specific mechanisms of audiovisual integration from brain oscillations. We used a novel methodology (speech-brain entrainment) and novel information theoretic measures (the Partial Information Decomposition) to quantify the interactions between audiovisual stimuli and dynamic brain signals.

Our methodology of speech-brain entrainment builds on recent studies suggesting that rhythmic components in brain activity that are temporally aligned to salient features in speech – most notably the syllable rate [5, 6, 13–15] – facilitate processing of both the auditory and visual speech inputs. The main advantage of speech-brain entrainment is that it replaces unspecific measures of activation with measures that directly quantify the coupling between the components of continuous speech (e.g. syllable rate) and frequency-specific brain activity, thereby tapping more directly into the brain mechanisms of speech segmentation and coding [14].

Our information theoretic measure of Partial Information Decomposition (PID; see Fig 1A and method for details) [16–18] addresses a perennial question in multisensory processing, the extent to which each sensory modality contributes uniquely to sensory representation in the brain vs. how different modalities (e.g. audio and visual) jointly contribute via a form of interaction. There are two main forms of interaction to consider. In redundant coding, the brain signal reflects common coding of auditory and visual speech. In synergistic coding the brain signal reflects a new representation of auditory and visual speech that is over and above representations based on each modality alone (i.e. a cross-modal enhancement of speech representation). Interestingly, both redundant and synergistic coding can be interpreted as reflecting different mechanisms of audiovisual information integration. The PID framework therefore provides a principled approach to investigate different cross-modal integration mechanisms (redundant and synergistic) in the human brain during naturalistic audiovisual speech processing. That is, to understand how neural representations of dynamic auditory and visual speech signals interact in the brain to form a unified perception.

**Fig 1.**
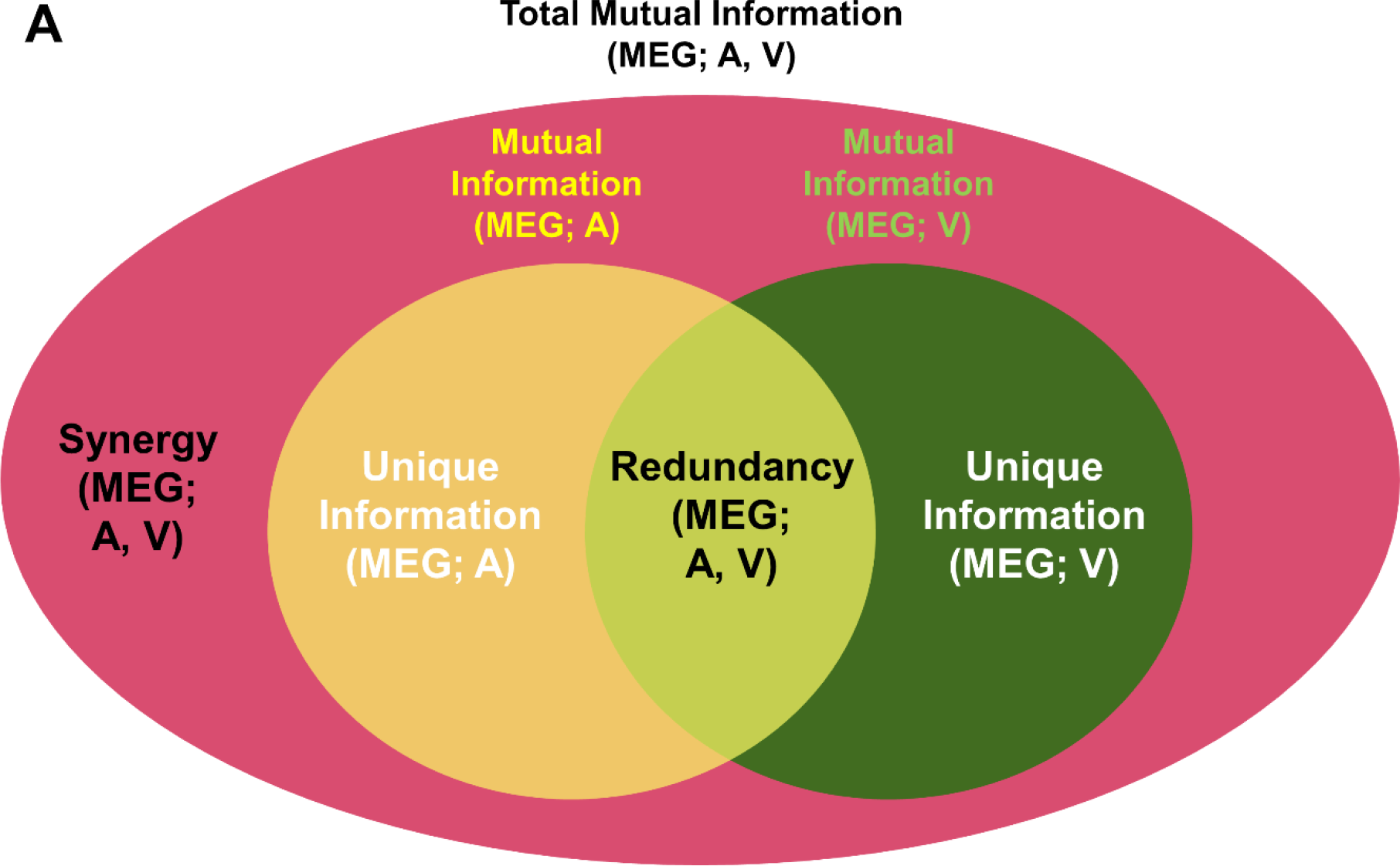
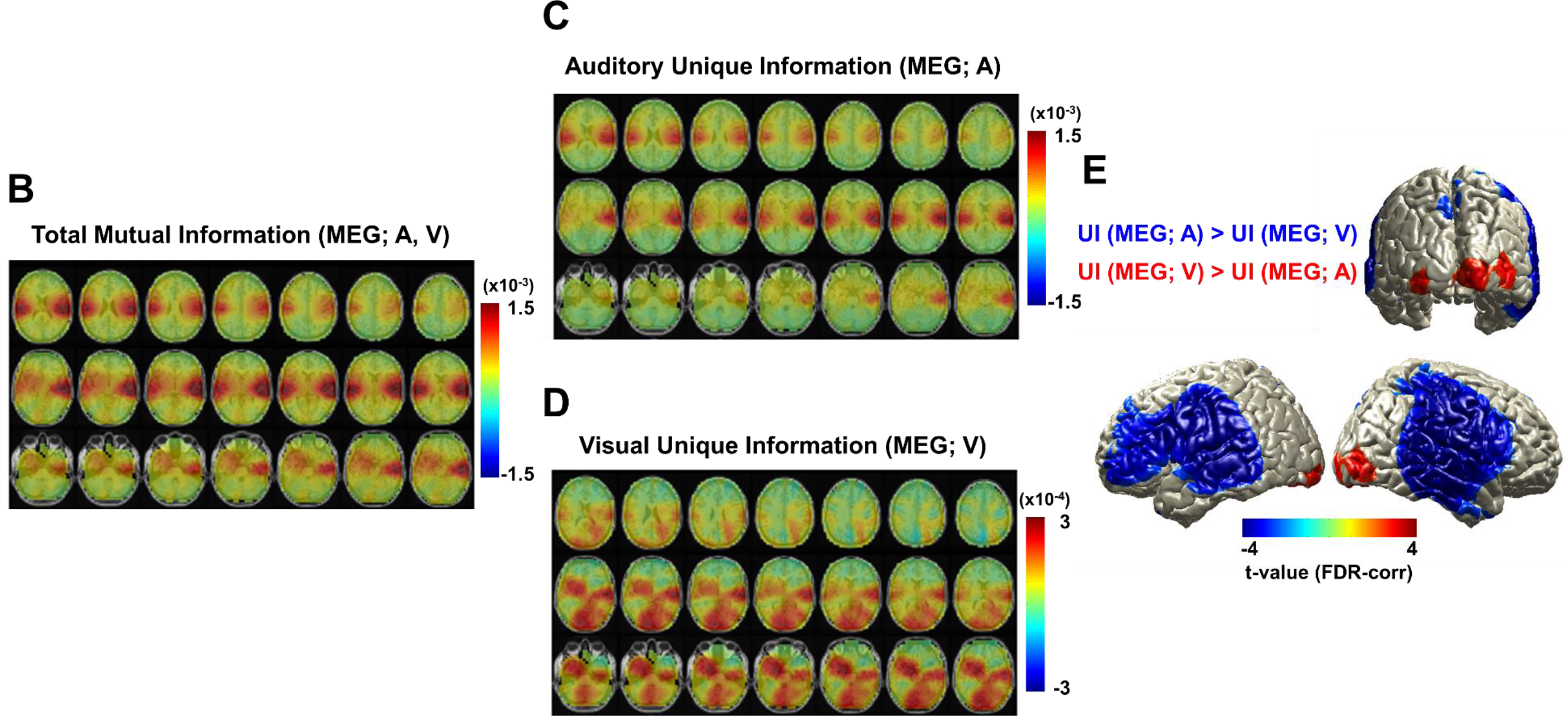
Partial Information Decomposition (PID) of audiovisual speech processing in the brain. A. Information structure of audiovisual multisensory inputs interacting with brain in which two source variables of auditory and visual speech signals (sound envelope and lip movement signal) hold about target variable (brain response). Ellipses indicate total mutual information (MEG; A,V), mutual information (MEG; A), mutual information (MEG; V) and the four distinct regions indicate unique information of auditory speech (MEG; A), unique information of visual speech (MEG; V), redundancy (MEG; A,V) and synergy (MEG; A,V), Figure modified from [16, 19]. See methods for details. B. Total mutual information quantified by PID was shown and this corresponds the ellipse of total mutual information (MEG; A, V) in A. C. Unique information of auditory speech (MEG; A) is shown in bilateral auditory cortex with stronger pattern in the right hemisphere. D. Unique information of visual speech (MEG; V) is observed in bilateral visual and auditory cortices. In order to observe overall patterns in each information map (B-D), we first normalized each information map by time-shifted surrogate map at individual level, then averaged across subjects. E. Unique information of visual speech and auditory speech were compared. Stronger unique information for auditory speech was found in bilateral auditory, temporal, inferior frontal areas and stronger unique information for visual speech was found in bilateral visual cortex (*P* < 0.05, false discovery rate (FDR) corrected).

Specifically, we recorded brain activity using MEG while participants attended to continuous audiovisual speech to entrain brain activity. We applied the PID to reveal where and how speech-entrained brain activity in different regions reflects different types of auditory and visual speech integration. In the first experimental condition, we used naturalistic audiovisual speech where attention to visual speech was not critical (‘All congruent’ condition). In the second condition, we added a second interfering auditory stimulus that was incongruent to the congruent audiovisual stimuli (‘AV congruent’ condition), requiring attention to visual speech to suppress the competing additional incongruent auditory input. In the third condition, both auditory stimuli were not congruent to visual stimulus (‘All incongruent’). This allows us to see how the congruence of audiovisual stimuli changes integration. We contrasted redundancy and synergy coding between the conditions to reveal differential effects of attention and congruence on multisensory integration mechanisms and behavioral performance.

## Results

We first studied partial information decomposition (PID) in an ‘All congruent’ condition (binaural presentation of speech with matching video) to understand multisensory representational interactions in the brain during processing of natural audiovisual speech. We used mutual information (MI) to quantify the overall dependence between the full multisensory dynamic stimulus time-course (broadband speech amplitude envelope and lip area for auditory and visual modalities respectively) and the recorded brain activity. This multisensory audiovisual MI (total mutual information (MEG; A, V), Fig 1B) includes unique unimodal as well as redundant and synergistic multisensory effects, which we can separate with the PID. The unique components are shown in Figs 1C-E. The total mutual information map (Fig 1B) shows multimodal stimulus entrainment in bilateral auditory/temporal areas and to lesser extent in visual cortex. Auditory unique information (MEG; A) is present in bilateral auditory areas (Fig 1C), where it accounts for the a large proportion of the total mutual information (MEG; A,V). Visual unique information (MEG; V) is present in both visual and auditory areas (Fig 1D), but overall visual entrainment is weaker than auditory entrainment (see S1 Fig for more details). Comparing the auditory unique information to visual unique information across subjects revealed stronger visual entrainment in bilateral visual cortex and stronger auditory entrainment in bilateral auditory, temporal, and inferior frontal areas (paired two-sided *t*-test, df: 43, *P* < 0.05, false discovery rate (FDR) corrected; Fig 1E).

To identify a frequency band where auditory and visual speech signals show significant dependencies, we computed MI between both signals and compared it to MI between non-matching auditory and visual speech signals for frequencies from 0 to 20 Hz. As expected, only matching audiovisual speech signals show significant MI peaking at 5 Hz (Fig 2A) consistent with previous results based on coherence (see Fig 2C in [5]). We therefore focus our further analysis on the 3-7 Hz frequency band that is known to correspond to the syllable rate in continuous speech [13] and within which amplitude envelope of speech is known to reliably entrain auditory brain activity [15, 20].

**Fig 2.**
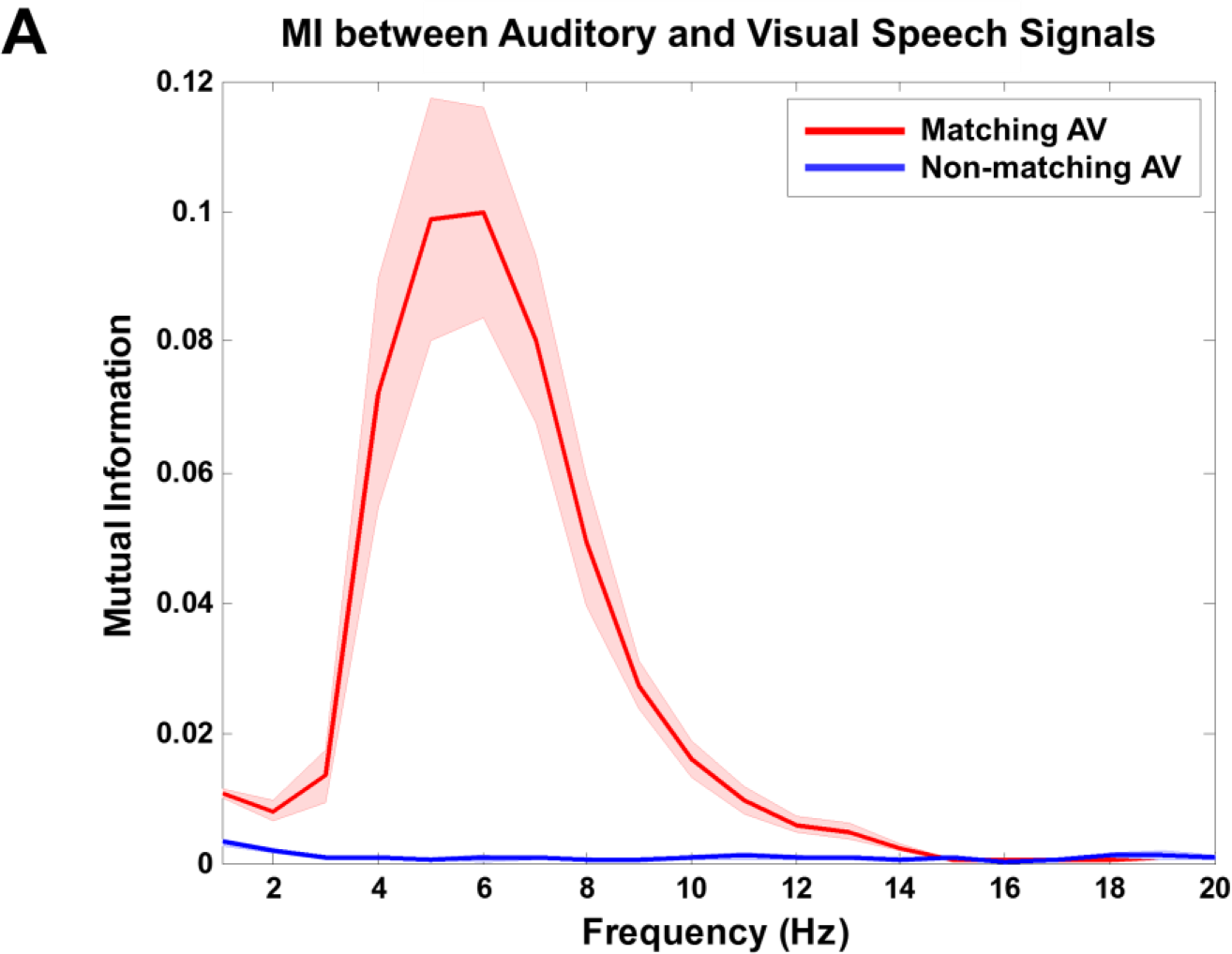
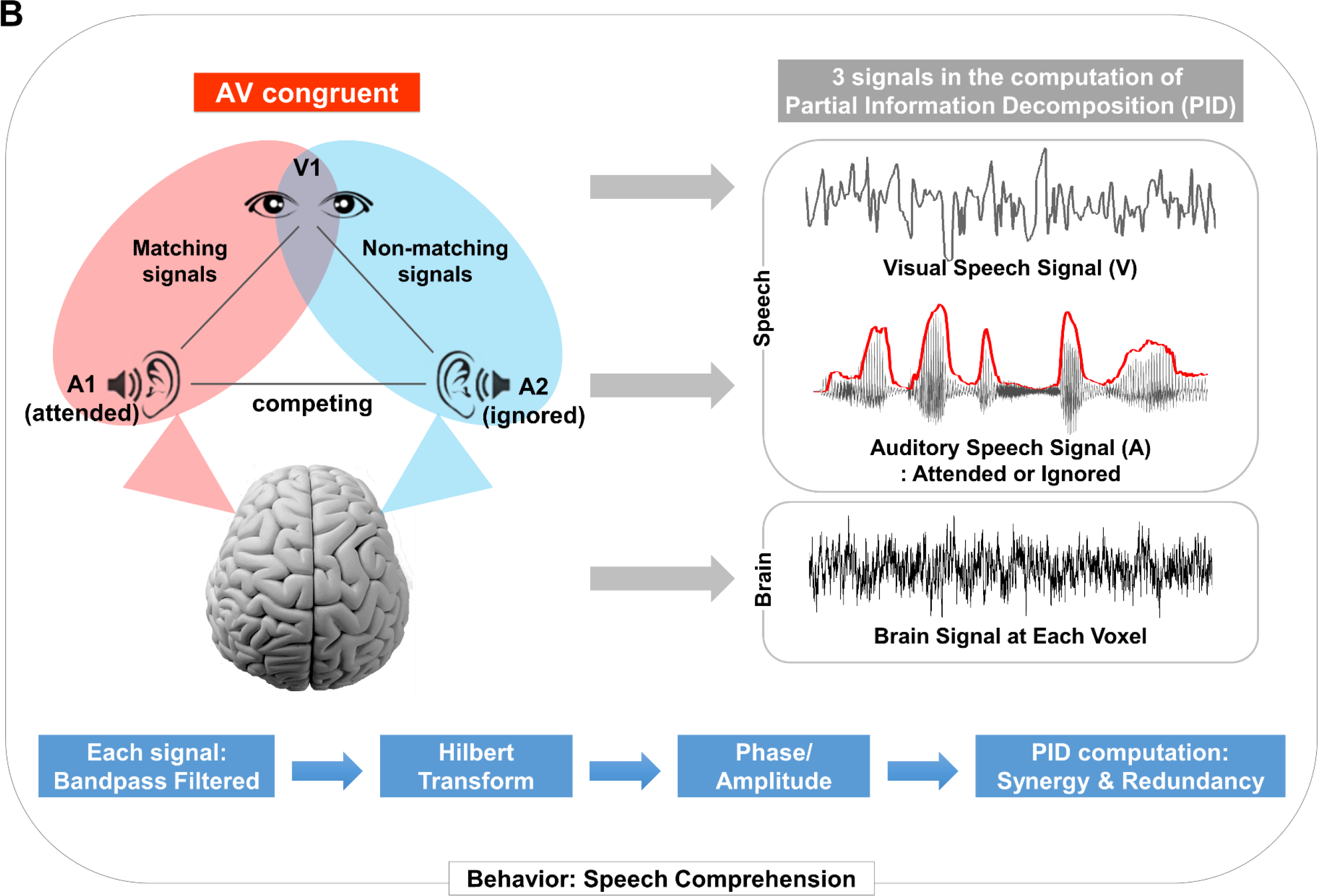
Mutual information between auditory and visual speech signals. A. To investigate Partial Information Decomposition (PID) in ‘AV congruent’ condition, first mutual information between auditory speech and visual speech signals was computed separately for matching and non-matching signals. Mutual information for matching auditory-visual speech signals shows nicely with a peak around 5 Hz (red line), whereas mutual information for non-matching signals is shown flat (blue line). B. Analysis of PID is shown for ‘AV congruent’ condition in which both matching and non-matching auditory-visual speech signals are present on the same brain response (MEG data). Two external speech signals (auditory speech envelope and lip movement signal) and brain signals were used in the PID computation. Each signal was bandpass filtered followed by Hilbert transform.

### Redundancy in left pSTG/S and synergy in left motor cortex

Next, we investigated how multimodal representational interactions are modulated by attention and congruence in continuous audiovisual speech. Here we focus on an ‘AV congruent’ condition where a congruent audiovisual stimulus pair is presented mon-aurally, together with an interfering non-matching auditory speech stimulus to the other ear (Fig 2B). This condition is of particular interest because visual speech (lip movement) is used to disambiguate the two competing auditory speech signals. Furthermore, it is ideally suited for our analysis because we can directly contrast representational interactions quantified with the PID in matching and non-matching audiovisual speech signals in the same data set (see Fig 2B).

Fig 3 shows corrected group statistics for the contrast of matching and non-matching audiovisual speech in the ‘AV congruent’ condition. Interestingly, redundant information is significantly stronger in left auditory and superior and middle temporal cortices (Fig 3A; Z-difference map at *P* < 0.005) for matching compared to non-matching audiovisual speech. In contrast, significantly higher synergistic information for matching compared to non-matching audiovisual speech is found in left motor and bilateral visual areas spreading along dorsal and ventral stream regions of speech processing [21] (Fig 3B; Z-difference map at *P* < 0.005). Next, we tested attention and congruence effects separately because the contrast of matching versus non-matching audiovisual speech confounds both effects.

**Fig 3.**
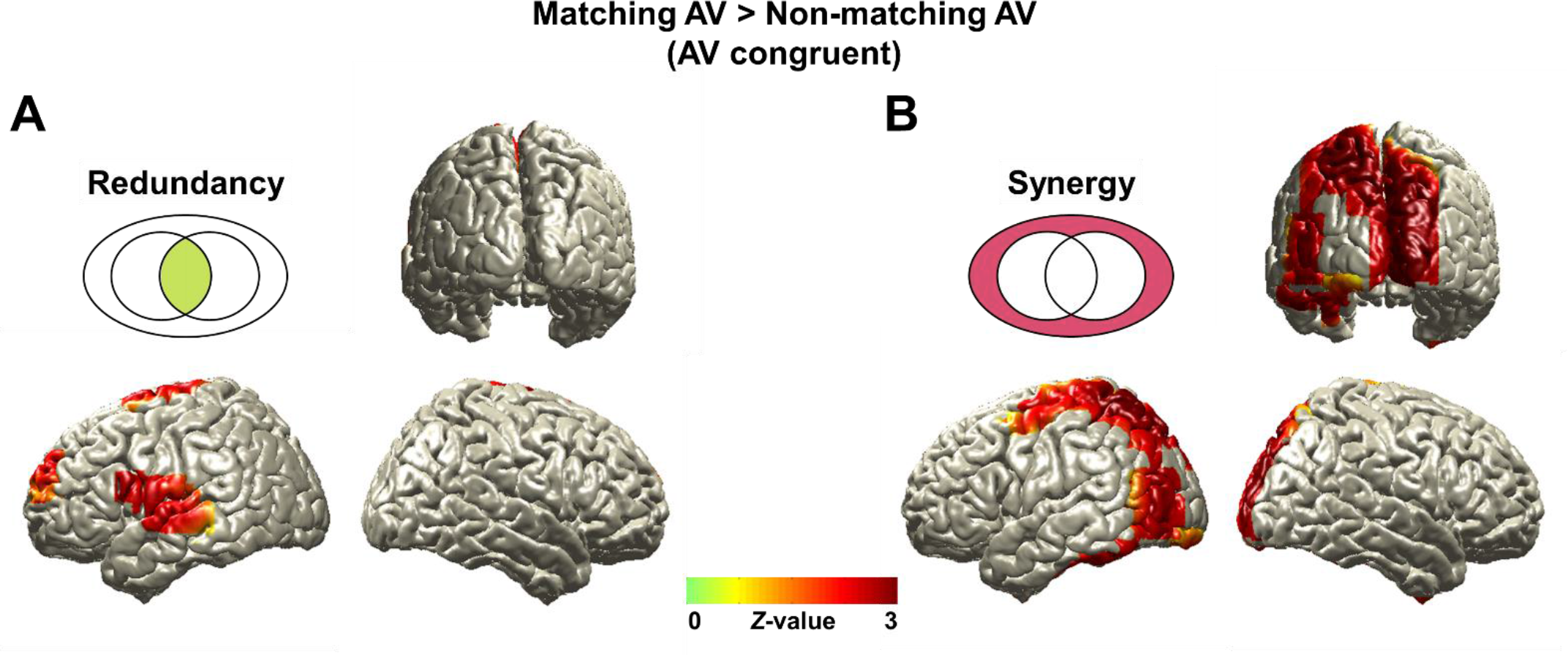
Redundancy and synergy revealed by Partial Information Decomposition (PID) for attention-modulated speech processing (‘AV congruent’ condition). Redundant and synergistic information of matching audiovisual speech signals in the brain compared to non-matching signals are shown. Each map (matching or non-matching in each information map) was firstly yielded to regression analysis using speech comprehension then transformed to standard Z maps and subtracted. A. Redundant information is localized in left auditory and superior and middle temporal cortices. B. Synergistic information is found in left motor and bilateral visual areas (Z-difference map at *P* < 0.005).

First, the congruence effect (‘AV congruent’ > ‘All incongruent’) shows higher redundant information in left inferior frontal region (BA 44/45) and posterior superior temporal gyrus and right posterior middle temporal cortex (Fig 4A; Z-difference map at *P* < 0.005) and higher synergistic information in superior part of motor/sensory cortices in left hemisphere (Fig 4B; Z-difference map at *P* < 0.005).

**Fig 4.**
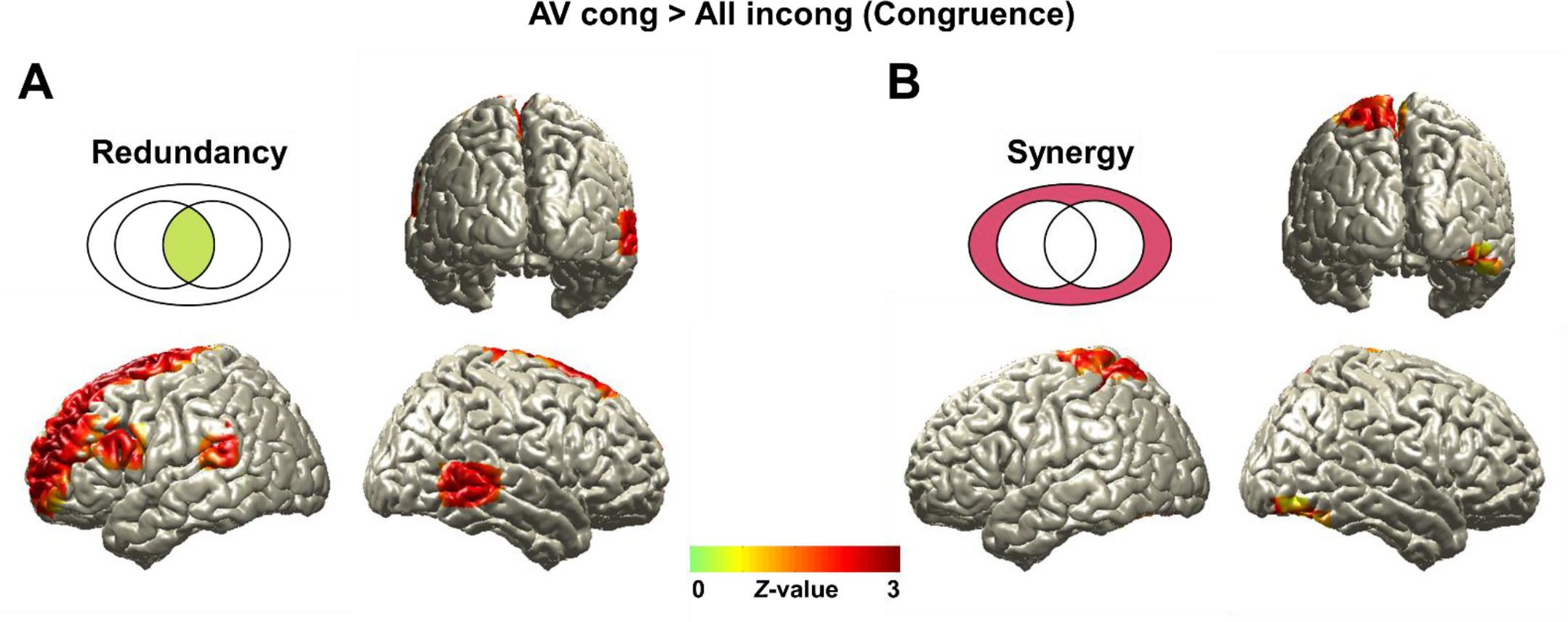
Redundancy and synergy in congruence effect. Comparison between conditions of matching versus non-matching audiovisual speech signals in ‘AV congruent’ condition entails both attention and congruence effects. To separate this effect, we additionally analyzed contrast for congruence (‘AV congruent’ > ‘All incongruent’) first. A. Redundancy for congruence effect is observed in left inferior frontal region and posterior superior temporal gyrus/sulcus (pSTG/S) and right posterior middle temporal cortex (*Z*-difference map at *P* < 0.005). B. Synergistic information for congruence effect is found in superior part of motor/sensory cortices in left hemisphere (*Z*-difference map at *P* < 0.005).

The attention effect (‘AV congruent’ > ‘All congruent’) shows higher redundant information in left auditory and temporal (superior and middle temporal cortices and posterior superior temporal gyrus/sulcus) areas and right inferior frontal and superior temporal cortex (Fig 5A; Z-difference map at *P* < 0.005). Higher synergistic information was localized in left motor cortex, inferior temporal cortex, and parieto-occipital areas (Fig 5B; Z-difference map at *P* < 0.005).

**Fig 5.**
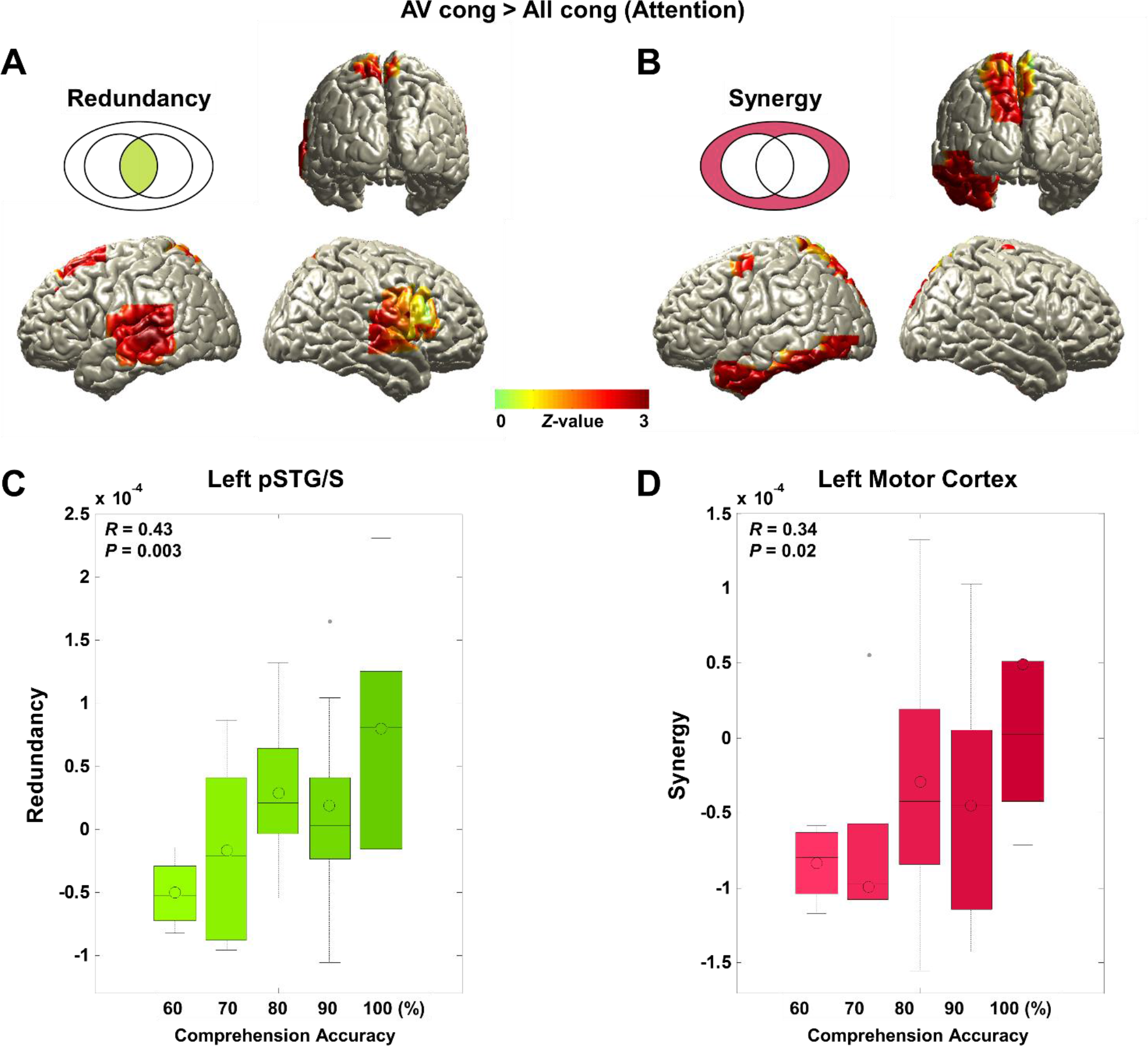
Redundancy and synergy in attention effect. Redundancy and synergy in attention (‘AV congruent’ > ‘All congruent’) are analyzed. Further to explore whether this effect is specific to ‘AV congruent’ condition (not because of decreased information in ‘All congruent’ condition), we extracted raw values of each information map at the local maximum voxel and correlated it with speech comprehension accuracy across subjects. A. Redundancy for attention effect was observed in left auditory and temporal (superior and middle temporal cortices and posterior superior temporal gyrus/sulcus) areas and right inferior frontal and superior temporal cortex (*Z*-difference map at *P* < 0.005). B. Synergistic information for attention effect was localized in left motor cortex, inferior temporal cortex, and parieto-occipital areas (*Z*-difference map at *P* < 0.005). C. Redundancy at the left posterior superior temporal region in ‘AV congruent’ condition was found to be positively correlated with speech comprehension accuracy (*R* = 0.43, *P* = 0.003). D. Synergy at the left motor cortex in ‘AV congruent’ condition was also positively correlated with speech comprehension accuracy across subjects (*R* = 0.34, *P* = 0.02). This finding suggests that redundant information in the left posterior superior temporal region and synergistic information in the left motor cortex in a challenging audiovisual speech condition support better speech comprehension.

In summary, theta-band activity in left posterior superior temporal gyrus/sulcus (pSTG/S) significantly represents redundant information about audiovisual speech more strongly in experimental conditions with higher attention and congruence. In contrast, synergistic information in the left motor cortex is more prominent in conditions requiring increased attention. Therefore, the increased relevance of visual speech in the ‘AV congruent’ condition leads to differential increase of different integration mechanisms in different brain areas – namely, increased redundancy in left pSTG/S and increased synergy in left motor cortex. For detailed local maps of interaction between predictors (auditory and visual speech signals) and target (MEG response) see S3 Fig.

### Performance scales with redundancy in left pSTG/S and synergy in left motor cortex

Next we investigated if the differential pattern of redundancy and synergy is of behavioral relevance in our most important condition – ‘AV congruent’ – where visual speech is particularly informative. To this end, we extracted raw values of redundancy for the location showing strongest redundancy in the left pSTG/S (Fig 5A) and synergy for the location showing strongest synergy in the left motor cortex (Fig 5B) for ‘AV congruent’ condition. After normalization with surrogate data (see Methods section), we computed correlation with performance measures (comprehension accuracy) across participants. Both redundancy in left pSTG/S (*R* = 0.43, *P* = 003; Fig 5C) and synergy in left motor cortex (*R* = 0.34, *P* = 0.02; Fig 5D) are significantly correlated with comprehension accuracy. These results suggest that the redundancy in left pSTG/S and synergy in left motor cortex under challenging conditions (i.e. in the presence of distracting speech) are related to perceptual mechanisms underlying comprehension.

## Discussion

In this study, we investigated how multisensory audiovisual speech rhythms are represented in the brain and how they are integrated for speech comprehension. We propose to study multisensory integration using information theory for the following reasons: First, it is a principled way to quantify the interactions between different stimuli. Second, interactions can be measured directly without resorting to statistical contrasts between conditions (e.g. AV > A+V). Third, information theory affords measures of interaction that cannot be computed in other ways (particularly specifying unique information and synergy) (Fig 1A) [16, 19]. The partial information decomposition (PID) allowed us to break down the relationships between the representations of auditory and visual speech inputs in the brain. We found that left posterior superior temporal region conveys speech information common to both auditory and visual modalities (redundancy) while left motor cortex conveys information that is greater than the linear summation of individual information (synergy). These results are obtained from low-frequency theta rhythm (3-7 Hz) signals, corresponding to syllable rate in speech components. Importantly, redundancy in pSTG/S and synergy in left motor cortex predict behavioral performance – speech comprehension accuracy – across participants.

A critical hallmark of multisensory integration in general, and audiovisual integration in particular, is the behavioral advantage conveyed by both stimulus modalities as compared to each single modality. Here we have shown that this process relies on at least two different mechanisms in two different brain areas, reflected in different representational interaction profiles revealed with synergy and redundancy.

### What do redundancy and synergy mean? Linking to audiovisual integration in fMRI studies

In fMRI studies, audiovisual speech integration has been studied by manipulating experimental conditions (e.g. [11, 22]). Changes in BOLD responses elicited by congruent audiovisual stimuli (AV) were compared to auditory-only (AO) or visual-only (VO), their sum (AO + VO) or their conjunction (AO ∩ VO). Greater activation for audiovisual condition (AV) compared to others were interpreted as audiovisual speech integration. Comparison to auditory-only (AO) or visual-only (VO) condition was regarded as less conservative criterion for integration and comparison to their summation (AO + VO) was considered as integration effect with supra-additivity. However, these contrasts are potentially confounded by condition-specific differences in attention, stimulus complexity, individual preference to individual stimulus modality (e.g. auditory over visual) and others. Similarly, this approach cannot identify how the brain makes use of the similar or complementary aspects of information in auditory and visual inputs.

Importantly, the PID can quantify the representational interactions between multiple sensory signals and the associated brain response in a single experimental condition where both sensory modalities are simultaneously present. In the PID framework, the unique contributions of a single (e.g. auditory) sensory modality to brain activity is directly quantified instead of relying on the statistical contrast between different conditions. Furthermore, the PID method allows the quantification of both redundant and synergistic interactions. In the context of audiovisual integration both types of interaction can be seen as integration effects. Redundant information refers to quantification of overlapping information content of the predictor variables (auditory and visual speech signals) and synergistic information refers to additional information gained from simultaneous observation of two predictor variables compared to observation of one. If we force a comparison between activation and information we could argue that redundancy is more related to the conjunction of each modality (AV ≈ AO ∩ VO) and synergy more related to supra-additivity (AV > (AO + VO)). However, it is important to keep in mind that activation and information are conceptually and computationally very different.

### Left posterior superior temporal region extracts common features from auditory and visual speech rhythms

Posterior superior temporal region (pSTG/S) has been implicated in audiovisual speech integration area by functional [23–25] and anatomical [26] neuroimaging. A typical finding in fMRI studies is that pSTG/S shows stronger activation for audiovisual (AV) compared to auditory-only (AO) and/or visual-only (VO) conditions. This was confirmed by a combined fMRI-TMS study in which the likelihood of McGurk fusion was reduced when TMS was applied individually to fMRI-localized pSTS, suggesting a critical role of pSTS in auditory-visual integration [12].

The redundant information in the same left superior temporal region in this study matches this notion that this region processes shared information from both modalities. We found this region not only in the congruence effect (‘AV congruent’ > ‘All incongruent’; Fig 4A) but also in the attention effect (‘AV congruent’ > ‘All congruent’; Fig 5A).

### Left motor cortex activity reflects synergistic information in audiovisual speech processing

Interestingly, we found the left motor cortex shows increased synergy for the matching vs non-matching audio stimuli of ‘AV congruent’ condition (Fig 3B). However, further analysis optimized for effects of attention and congruence revealed two slightly different precentral areas – with the area that shows strongest synergy change with attention (Fig 5B) located more lateral and anterior compared to the area identified in the congruence (Fig 4B). Interestingly, the motor region in the attention contrast is consistent with the area in our previous study that showed entrainment to lip movements during continuous speech that correlated with speech comprehension [5]. In another study we identified this area as the source of top-down modulation of activity in the left auditory cortex [20]. The definition of synergistic information in our context refers to more information gained from the simultaneous observation of auditory and visual speech compared to the observation of each alone. When it comes to the attention effect (‘AV congruent’ > ‘All congruent’), ‘AV congruent’ condition requires paying more attention to auditory and visual speech than the ‘All congruent’ condition does, even though the speech signals to be attended match the visual stimulus in both conditions. Thus, this synergy effect in the left motor cortex can be explained by a net attention effect at the same level of stimulus congruence. This effect is likely driven by stronger attention to visual speech which is informative for the disambiguation of the two competing auditory speech streams [5]. This notion is plausible because it is supported by directional information analysis which shows that the left motor cortex better predicts upcoming visual speech in the ‘AV congruent’ condition where attention to visual speech is crucial (S2 Fig B,D).

In summary, we demonstrate how information theoretic tools can provide a new perspective on audiovisual integration, by explicitly quantifying both redundant and synergistic cross-modal representational interactions. This reveals two distinct profiles of audiovisual integration, that are supported by different brain areas (left motor cortex and left pSTG/S) and are differentially recruited under different listening conditions.

## Materials and Methods

### Participants

Data from 44 subjects were analyzed (26 females; age range: 18-30 years; mean age: 20.54 ± 2.58 years). Another analysis of these data was presented in a previous report [5]. All subjects were healthy, right-handed and had normal or corrected-to-normal vision and normal hearing. None of the participants had a history of developmental, psychological, or neurological disorders. They all provided informed written consent before the experiment and received monetary compensation for their participation. The study was approved by the local ethics committee (CSE01321; College of Science and Engineering, University of Glasgow) and conducted in accordance with the ethical guidelines in the Declaration of Helsinki.

### Stimuli and Experiment

We used audiovisual video clips of a professional male speaker talking continuously (7-9 minutes) which were used in our previous study [5]. The talks were originally taken from TED talks (www.ted.com/talks/) and edited to be appropriate to the stimuli we used (e.g. editing words referring to visual materials, the gender of the speaker etc.).

High-quality audiovisual video clips were filmed by a professional filming company with sampling rate of 48 kHz for audio and 25 fps (frame per second) for video in 1920 x 1080 pixels.

In order to validate stimuli, eleven videos were rated by 33 participants (19 females; aged 18-31 years; mean age: 22.27 ± 2.64 years) in terms of arousal, familiarity, valence, complexity, significance (informativeness), agreement (persuasiveness), concreteness, self-relatedness, and level of understanding using Likert scale [27] 1 to 5 (for an example of concreteness, 1: very abstract, 2: abstract, 3: neither abstract nor concrete, 4: concrete, 5: very concrete). Eight talks were finally selected for the MEG experiment by excluding talks with mean scores of 1 and 5.

Questionnaires for each talk were validated in a separate behavioral study (16 subjects; 13 females; aged 18-23 years; mean age: 19.88 ± 1.71 years). These questionnaires are designed to assess the level of speech comprehension. Each questionnaire consists of 10 questions about a given talk to test general comprehension (e.g., “*What is the speaker’s job*?”) and were validated in terms of accuracy (the same level of difficulty), response time, and the length (word count).

Experimental conditions used in this study were ‘All congruent’, ‘All incongruent’, ‘AV congruent’. In each condition (7-9 min), one video recording was presented and two (matching or non-matching) auditory recordings were presented to the left and the right ear, respectively. The ‘All congruent’ condition is a natural audiovisual speech condition where auditory stimuli to both ears and visual stimuli are congruent. The ‘All incongruent’ condition is where all three stimuli are from different videos and participants are instructed to attend to auditory information presented to one ear. The ‘AV congruent’ condition is where only one of auditory stimuli matches the visual information, and the speech presented to the other ear serves as a distraction. Participants attend to the talk that matches visual information. Participants were instructed to fixate on the speaker’s lip all the time in all experimental conditions. In ‘All congruent’ condition (natural audiovisual speech), they were instructed to ignore the color of the fixation cross and just to attend to both sides naturally.

A fixation cross (either yellow or blue color) was overlaid on the speaker’s lip to indicate the auditory stimulus to pay attention to (left or right ear, e.g. “If the color of fixation cross is yellow, please attend to left ear speech”). The color was counterbalanced across subjects. For the recombination and editing of audiovisual talks we used Final Cut Pro X (Apple Inc., Cupertino, CA).

Half of the 44 participants attended to speech in the left ear and the other half attended to speech in the right ear. There was no significant difference in comprehension accuracy between groups (two sample *t*-test, df: 42, *P* > 0.05). In this study, we pooled across both groups for data analysis.

The stimuli were presented with Psychtoolbox [28] in MATLAB (MathWorks, Natick, MA). Visual stimuli were delivered with a resolution of 1280 × 720 pixels at 25 fps (mp4 format). Auditory stimuli were delivered at 48 kHz sampling rate via a sound pressure transducer through two 5 meter-long plastic tubes terminating in plastic insert earpieces.

A comprehension questionnaire was administered about the attended speech separately for each condition.

## Data acquisition

Cortical neuromagnetic signals were recorded using a 248-magnetometers whole-head MEG (Magnetoencephalography) system (MAGNES 3600 WH, 4-D Neuroimaging) in a magnetically shielded room. The MEG signals were sampled at 1,017 Hz and were denoised with information from the reference sensors using the denoise_pca function in FieldTrip toolbox [29]. Bad sensors were excluded by visual inspection and electrooculographic (EOG) and electrocardiographic (ECG) artifacts were eliminated using independent component analysis (ICA). An eye tracker (EyeLink 1000, SR Research Ltd.) was used to examine participants’ eye gaze and movements, to ensure that they fixated on the speaker’s lip movements.

Structural T1-weighted magnetic resonance images (MRI) of each participant were acquired at 3 T Siemens Trio Tim scanner (Siemens, Erlangen, Germany) with the following parameters: 1.0 × 1. 0 × 1.0 mm^3^ voxels; 192 sagittal slices; Field of view (FOV): 256 × 256 matrix.

## Data analysis

Information theoretic quantities were estimated with the Gaussian-Copula Mutual Information (GCMI) method [30] (https://github.com/robince/gcmi). Partial information decomposition (PID) analysis was performed with the GCMI approach in combination with an open source PID implementation in MATLAB, which implements the PID [17, 18] with a redundancy measure based on common change in local surprisal [16] (https://github.com/robince/partial-info-decomp). For statistics and visualization, we used the FieldTrip Toolbox [29] and in-house MATLAB codes. We followed the suggested guidelines [31] for MEG studies.

**MEG-MRI co-registration.** Structural MR images of each participant were co-registered to the MEG coordinate system using a semi-automatic procedure. Anatomical landmarks (nasion, bilateral pre-auricular points) were identified before the MEG recording and also manually identified in the individual’s MR images. Based on these landmarks, both MEG and MRI coordinate systems were initially aligned. Subsequently, numerical optimization was achieved by using the ICP algorithm [32].

**Source localization.** A head model was created for each individual from their structural MRI using normalization and segmentation routines in FieldTrip and SPM8. Leadfield computation was performed based on a single shell volume conductor model [33] using a 8-mm grid defined on the template provided by MNI (Montreal Neurological Institute). The template grid was linearly transformed into individual head space for spatial normalization. Cross-spectral density matrices were computed using Fast Fourier Transform on 1-s segments of data after applying multitaper. Source localization was performed using DICS beamforming algorithm [34] and beamformer coefficients were computed.

**Auditory speech signal processing.** The amplitude envelope of auditory speech signals was computed following the approach reported in [35]. We constructed eight frequency bands in the range 100-10,000 Hz to be equidistant on the cochlear map [36]. The auditory sound speech signals were band-pass filtered in these bands using a fourth-order forward and reverse Butterworth filter. Then Hilbert transform was applied to obtain amplitude envelopes for each band of signal. These signals were then averaged across bands and resulted in a wideband amplitude envelope. For further analysis, signals were downsampled to 250 Hz.

**Visual speech signal processing.** A lip movement signal was computed using an in-house Matlab script. We first extracted the outline lip contour of the speaker for each frame of the movie stimuli. From the lip contour outline we computed the frame-by-frame lip area (area within lip contour). This signal was resampled at 250 Hz to match the sampling rate of the preprocessed MEG signal and auditory sound envelope signal. We reported the first demonstration of visual speech entrainment using this lip movement signal [5].

### Estimating mutual information (MI) and other information theoretic quantities

**Shannon’s Information Theory** [37]. Information theory was originally developed to study man-made communication systems, however it also provides a theoretical framework for practical statistical analysis. It has become popular for the analysis of complex systems in a range of fields, and has been successfully applied in neuroscience to spike trains [38, 39], LFPs [40, 41], EEG [42, 43], MEG time-series data [15, 20]. Mutual information is a measure of statistical dependence between two variables, with a meaningful effect size measured in bits (see [30] for a review). MI of 1 bit corresponds to a reduction of uncertainty about one variable of a factor 2 after observation of another variable. Here we estimate MI and other quantities using Gaussian-Copula Mutual Information (GCMI) [30]. This provides a robust semi-parametric lower bound estimator of mutual information, by combining the statistical theory of copulas with the closed form solution for the entropy of Gaussian variables. Crucially, this method performs well for higher dimensional responses as required for measuring three-way statistical interactions (see below) and allows estimation over circular variables like phase.

### Mutual information (MI) between auditory and visual speech signals

Following the GCMI method [30], we normalized the complex spectrum by its amplitude to obtain a 2d representation of the phase as points lying on the unit circle. We then rank-normalized the real and imaginary parts of this normalized spectrum separately, and used the multivariate GCMI estimator to quantify the dependence between these two 2d signals. This gives a lower bound estimate of the MI between the phases of the two signals.

To determine the frequency of interest for the main analysis (partial information decomposition; PID), we computed MI between auditory (A) and visual (V) speech signals for the matching AV and non-matching AV signals from all the stimuli we used. As shown in Fig 2A, there was no relationship between non-matching auditory and visual stimuli, but there was a frequency dependent relationship for matching stimuli peaking in the band 3-7 Hz. This is consistent with previous results using coherence measures [5, 35]. This frequency band corresponds to the syllable rate and is known to show robust phase coupling between speech and brain signals.

### Partial Information Decomposition (PID) theory

We seek to study the relationships between the neural representations of auditory (here amplitude envelope) and visual (here dynamic lip area) stimuli during natural speech. Mutual information can quantify entrainment of the MEG signal by either or both of these stimuli, but cannot address the relationship between the two entrained representations - their representational interactions. The existence of significant auditory entrainment revealed with MI demonstrates that an observer who saw a section of auditory stimulus would be able to, on average, make some prediction about the MEG activity recorded after presentation of that stimulus (this is precisely what is quantified by MI, without the need for an explicit model). Visual MI reveals the same for the lip area. However, a natural question is then whether these two stimulus modalities provide the same information about the MEG, or provide different information. If an observer saw the auditory stimulus, and made a corresponding prediction for the MEG activity, would that prediction be improved by observation of the concurrent visual stimulus, or would all the information about the likely MEG response available in the visual stimulus already be available from the related auditory stimulus? Alternatively, would an observer who saw both modalities together perhaps be able to make a better prediction of the MEG, on average, then would be possible if the modalities were not observed simultaneously?

This is conceptually the same question that is addressed with techniques such as Representational Similarity Analysis (RSA [44] or cross-decoding [45]. RSA determines similar representations by comparing the pairwise similarity structure in responses evoked by a stimulus set usually consisting of many examplars with hierarchical categorical structure. If the pattern of pairwise relationships between stimulus evoked responses is similar between two brain areas, it indicates there is a similarity in how the stimulus ensemble is represented. Cross-decoding works by training a classification or regression algorithm in one experimental condition or time region, and then testing its performance in another experimental region or time region. If it performs above chance on the test set, this demonstrates some aspect of the representation in the data that the algorithm learned in the training phase, is preserved in the second situation. Both these techniques address the same conceptual issue of representational similarity, which is measured with redundancy in the information theoretic framework, but have specific experimental design constraints, and are usually used to compare different neural responses (recorded from different regions, time periods or with different experimental modalities). The information theoretic approach is more flexible, and can be applied both to simple binary experimental conditions, as well as continuous valued dynamic features extracted from complex naturalistic stimuli such as those we consider here. Further, it allows us to study representational interactions between stimulus features (not only neural responses), and provides the ability to quantify synergistic as well as redundant interactions.

We can address this question with information theory through a quantity called ‘Interaction Information’ [16, 46], which is defined as follows:

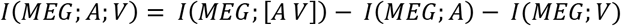

This quantifies the difference between the MI when the two modalities are observed together and the sum of the MI when each modality is considered alone. If the interaction information is positive, this indicates a synergistic representation. The two stimuli provide a better prediction when they are considered together than would be expected from observing each individually. If the interaction information is negative this indicates a redundant, or shared representation. Some of what is learnt about the neural response from the visual stimulus is already obtained from observation of the auditory stimulus.

Interaction information is the difference between synergy and redundancy [17] and therefore measures a net effect. It is possible to have zero interaction information even in the presence of strong redundant and synergistic interactions (for example over different ranges of the stimulus space) that cancel out in the net value. The methodological problem of fully separating redundancy and synergy has recently been addressed with the development of a framework called the Partial Information Decomposition (PID) [17–19, 47]. This provides a mathematical framework to obtain decomposition of mutual information into unique, redundant and synergistic components. The PID requires a measure of information redundancy. Here we use recently proposed measure of redundancy based on pointwise common change in surprisal; Iccs [16]. This approach starts from interaction information, but breaks down the conflated redundant and synergistic effects by only counting pointwise terms in the interaction information calculation that unambiguously correspond to a redundant interaction. This is the only redundancy measure which corresponds to an intuitive notion of overlapping information content and is defined for more than 2 variables and for continuous systems. We use it here in a continuous Gaussian formulation together with the rank-normalization approach of GCMI.

The PID allows us to separate the redundant and synergistic contributions to the interaction information, as well as the unique information in each modality (Fig 1). For clarity, we restate the interpretation of these terms in this experimental context.

**Unique Information (MEG; A)** – This quantifies that part of the MEG activity than can be explained or predicted only from the auditory speech envelope. This necessarily means it represents entrainment to speech envelope features that are not common to the lip movement (i.e., the relationships quantified in Fig 1C).

**Unique Information (MEG; V)** – This quantifies that part of the MEG activity that can be explained or predicted only from the visual lip area. This necessarily means it represents entrainment to speech envelope features that are not common to the speech envelope (i.e., the relationships quantified in Fig 1D).

**Redundancy (MEG; A V)** – This quantifies the information about the MEG signal that is common to or shared between the two modalities. Alternatively, this quantifies the representation in the MEG of the variations that are common to both signals.

**Synergy (MEG; A V)** – This quantifies the extra information that arises when both modalities are considered together. It indicates that prediction of the MEG response is improved by considering the dynamic relationship between the two stimuli, over and above what could be obtained from considering them individually.

### Partial Information Decomposition (PID) analysis

For brain signals, frequency-specific brain activation time-series were computed by applying the beamformer coefficients to the MEG data filtered in the same frequency band (fourth order Butterworth filter, forward and reverse, center frequency ± 2 Hz). The auditory and visual speech signals were filtered in the same frequency band. MEG signals were shifted by 100 ms as in previous studies [5, 15] to compensate for delays between stimulus presentation and cortical responses. Then, each map of PID was computed using these auditory, visual speech signals and source-localized brain signal for each voxel and each frequency band across 1-s-long data segments overlapping by 0.5 s.

As described above (MI between auditory and visual speech signals) the complex spectra obtained from the Hilbert transform were amplitude normalized, and the real and imaginary parts were each rank-normalized. The covariance matrix of the full 6-dimensional signal space was then computed which completely describes the Gaussian-Copula dependence between the variables. The PID was applied with redundancy measured by pointwise common change in surprisal (Iccs) [16] for Gaussian variables.

This calculation was performed independently for each voxel, resulting in volumetric maps for the four PID terms (redundant information, unique information of auditory speech, unique information of visual speech, synergistic information) for each frequency band in each individual. This computation was performed for all experimental conditions: ‘All congruent’, ‘All incongruent’, ‘AV congruent’.

In addition, surrogate maps were created by computing the same decomposed information maps between brain signals and time-shifted speech signals for each of the four experimental conditions in each individual. Visual speech signals were shifted for 30 s and auditory speech signals were shifted for 60 s. This surrogate data provides an estimate of each information map that can be expected by chance for each condition. This surrogate data is not used to create a null distribution but to estimate analysis bias. The surrogate data is used in analysis for Figs 1B-D, 5C-D, and S1, S2, S4 Figs.

### Delayed Mutual Information analysis

We used Delayed Mutual Information to investigate to what extent brain areas predict upcoming auditory or visual speech. Delayed mutual information refers to mutual information between two signals offset with different delays. If there is significant MI between brain activity at one time, and the speech signal at a later time, this shows that brain activity contains information about the future of the speech signal. We calculated delayed MI between each voxel and the two speech stimuli, from 0 ms to 500 ms with a 20 ms step (S2 Fig). Directed Information or Transfer Entropy [48, 49], is based on the same principle but additionally conditions out the past of the speech signal, to ensure the delayed interaction is providing new information over and above that available in the past of the stimulus. Here, since the delayed MI peaks are clear and well isolated from the 0 lag we present the simpler measure, but transfer entropy calculations revealed similar effects (results not shown).

### Selection of brain regions for Partial Information Decomposition (PID) analyses predictive of speech

We selected eight brain regions to test a predictive mechanism, i.e., does MEG activity predict upcoming speech. Regions were selected from the maximum coordinates of contrast for attention and congruence effects shown in Figs 4 and 5. The abbreviation used in the nodes in the Figures, MNI coordinates, Talairach coordinates [50] and Brodmann area (BA) are shown in the parenthesis: Left auditory cortex (A1; MNI = [−36 −24 8]; TAL = [−35.6 −22.9 8.5]; BA 41/22), left visual cortex (V1; MNI = [−28 −88 −8]; TAL = [−27.7 −85.6 −2.5]; BA 18), left posterior superior temporal gyrus (pSTG; MNI = [−60 −24 0]; TAL = [−59.4 −23.3 1.2]; BA 21/22), left motor cortex (M1; MNI = [−44 0 64]; TAL = [−43.6 2.9 58.8]; BA 6), left supplementary motor area (SMA; MNI = [−4 0 48]; TAL = [−4.0 2.2 44.1]; BA 6), left inferior frontal gyrus (IFG; MNI = [−64 16 24]; TAL = [-63.4 16.6 21.3]; BA 44/45), right inferior frontal gyrus (IFG; MNI = [60 8 16]; TAL = [59.4 8.5 14.3]; BA 44), left precuneus (Prec; MNI = [−4 −72 64]; TAL = [−4 −66.8 62.3]; BA 7).

### Partial Information Decomposition (PID) analysis predictive of speech

The PID analysis described above was computed to investigate cross-modal AV representational interactions in an individual brain region. But both PID and interaction information can be applied also to consider representational interactions between two brain regions to a single stimulus feature (as RSA is normally applied). To understand representational interactions between brain regions predictive of speech signals, we computed PID values with activity from two brain regions as the predictor variables and a unimodal speech signal (auditory or visual) as a target variable. For this, we used the eight brain regions (see above) and computed the PID for each pair of eight regions predictive of visual speech or auditory speech. This resulted in 28 pairwise computations (n(n-1)/2).

Here, redundant information between two brain regions means they both provide the same prediction of the upcoming speech signal. A synergistic interaction demonstrates that the particular dynamic relationship between the neural activity in the two regions is itself predictive of speech, in a way that the direct recorded MEG in each region alone is not (S4 Fig).

**Statistics.** Group statistics was performed on the data of all 44 participants in the FieldTrip. First, individual volumetric maps for each calculation (Mutual Information, Unique Information, Redundancy, Synergy) were smoothed with a 10-mm Gaussian kernel. Then they were subjected to dependent *t*-statistics using nonparametric randomization (Monte Carlo randomization) for comparisons between experimental conditions or to surrogate data. Results are reported after multiple comparison correction was performed using FDR (False Discovery Rate) [51].

For the maps relating synergistic and redundant PID relevant for behavior, each information map (unique, redundant, and synergistic map) was subjected to regression analysis. In the regression analysis, we detected brain regions that were positively correlated to comprehension accuracy using nonparametric randomization (Monte Carlo randomization). Then, regression *t*-maps were converted to standard *Z*-map (*Z*-transformation) and subtracted between conditions (*P* < 0.005).

## Author Contributions

The authors have made the following declarations about their contributions: Conceived and designed the experiments: HP, GT, JG. Performed the experiments: HP. Analyzed the data: HP. Contributed reagents/materials/analysis tools: HP, RI, JG. Wrote the paper: HP, RI, PGS, GT, JG.

## Acknowledgments

JG is supported by the Wellcome Trust (098433), GT is supported by the Wellcome Trust (098434), and PGS is supported by the Wellcome Trust (107802). The funders had no role in study design, data collection and analysis, decision to publish, or preparation of the manuscript.

## Supporting Information

**S1 Fig.**
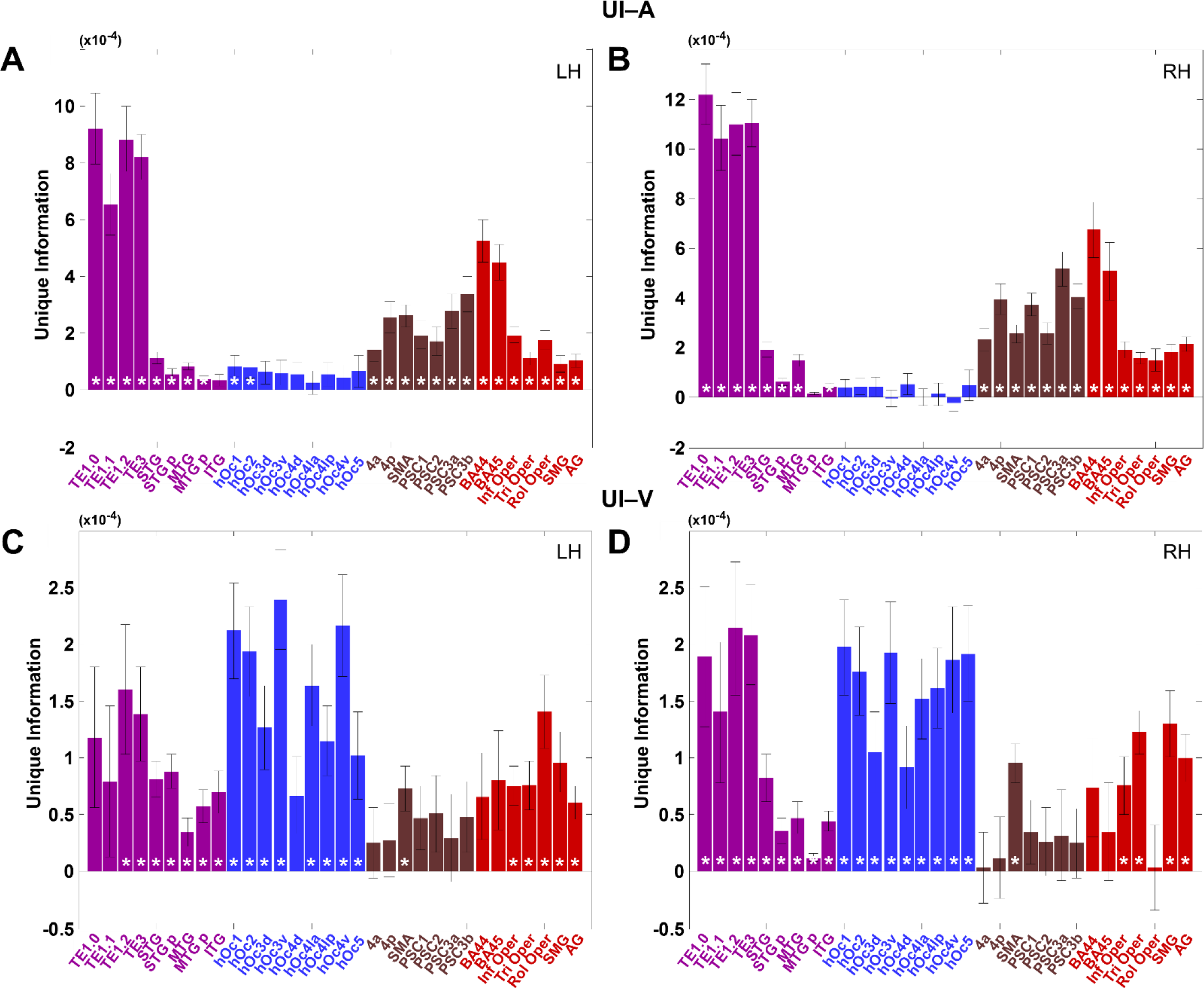
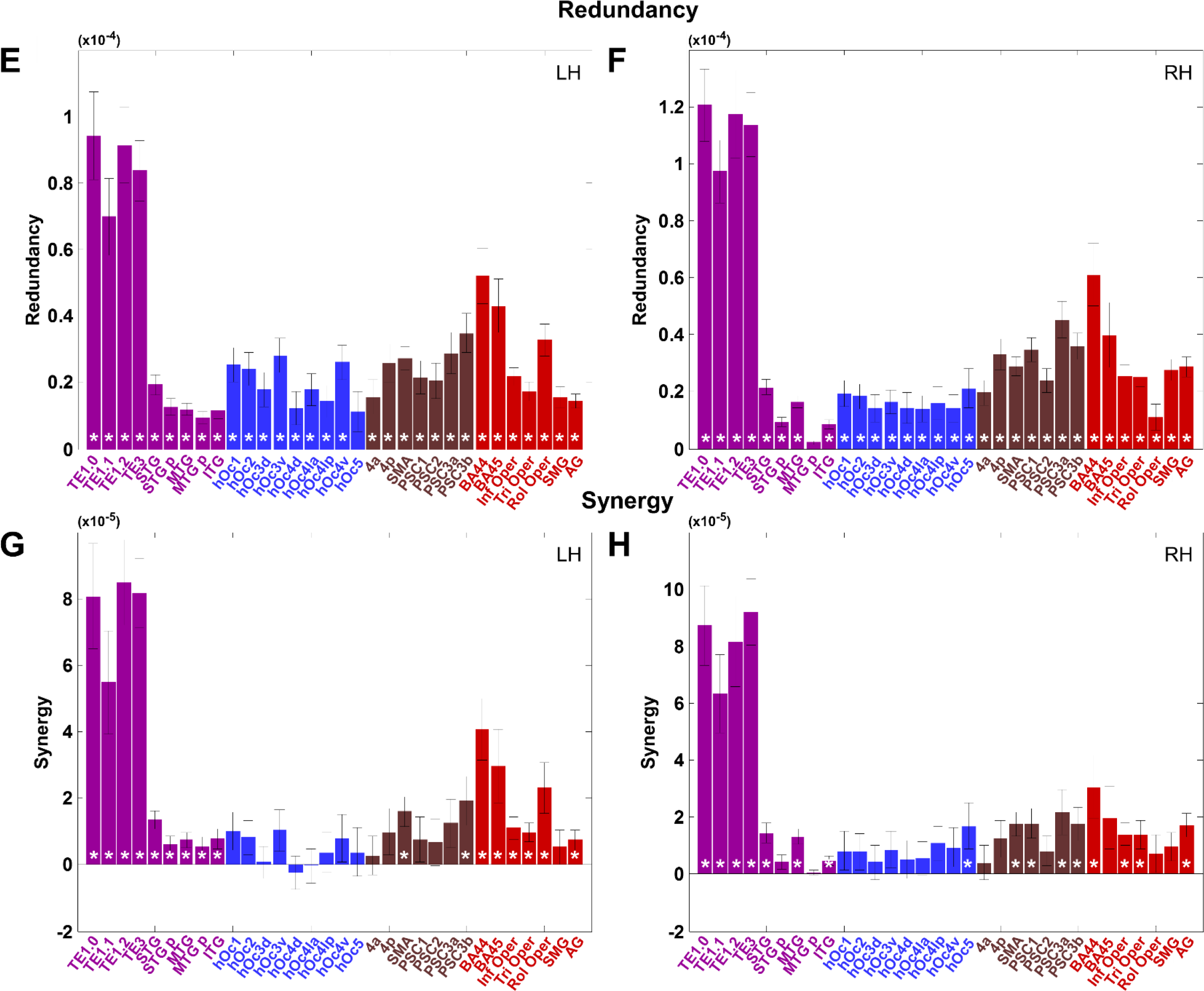
Neural decomposition of natural audiovisual speech (‘All congruent’ condition). In order to define characteristics of decomposed information in naturalistic audiovisual speech condition (All congruent), we used predefined ROI maps from SPM Anatomy Toolbox (version 2.1) [52] and Automated Anatomical Labeling (AAL) [53]. SPM Anatomy Toolbox provides probabilistic cytoarchitectonic maps which provides stereotaxic information on the location and variability of cortical areas in the MNI (Montreal Neurological Institute) space. AAL maps provide anatomical parcellation of the spatially normalized single-subject high-resolution T1 of MNI space. Both ROI toolbox provide complementary ROI maps to each other, so that we used both toolboxes. We were interested in each decomposed PID profile in natural speech condition in four main areas: Auditory/Temporal, Visual, Motor/Sensory, Language-related areas. Auditory/Temporal area (purple) includes primary auditory cortex (TE 1.0, TE 1.1, TE 1.2), higher auditory cortex (TE 3), superior temporal gyrus (STG), superior temporal pole (STG p), middle temporal gyrus (MTG), middle temporal pole (MTG p), inferior temporal gyrus (ITG). Visual area (blue) includes BA17/V1 (hOC1), BA18/V2 (hOC2), dorsal extrastriate cortex (hOC3d/hOC4d), ventral extrastriate cortex (hOC3v/hOC4v), lateral occipital cortex (hOc4la, hOc4lp), V5/MT+ area (hOc5). Motor/Sensory area (brown) includes Areas 4a and 4p, supplementary motor area (SMA), and primary somatosensory cortex Areas 1, 2, 3a, 3b. Language-related area (red) includes BA44, BA45, Inferior frontal opercular part (Inf Oper), Inferior frontal triangular part (Tri Oper), Rolandic operculum (Rol Oper), supramarginal gyrus (SMG), and angular gyrus (AG). We first transformed the dimension of each ROI map to the dimension of our source space data, then we extracted each information (unique unimodal information for auditory and visual speech, redundancy and synergy) of bandpass-filtered (low frequencies 1-7 Hz) phase data from each ROI and then each information value was averaged within the ROI. This was performed for All congruent condition and time-shifted surrogate data. Each information data was averaged across all subjects after subtracted by surrogate data within individual (mean ± s.e.m). Data shown per each hemisphere (LH, RH). Statistics compared to surrogate data was also performed and shown with asterisk in each bar when it is significant (paired two-sided t-test; *P* < 0.05). First for unimodal unique information (UI-A, UI-V), as expected, auditory unique information (A, B) showed stronger unique information in primary auditory cortices while visual unique information (C, D) showed stronger unique information in visual cortices as well as auditory areas suggesting auditory cortex might be involved in visual speech processing. Next, for Redundancy (E,F) and Synergy (G,H), overall redundant information in both hemispheres shows stronger than synergistic information in all main areas. This was expected considering the same audiovisual inputs in this condition (All congruent). In auditory/temporal areas, both redundant and synergistic information were significantly different from time-shifted surrogate data. This pattern is the same in both hemispheres. However, in visual areas, only redundant information is significant whereas nearly none of synergistic information in visual cortices remain significant. In motor/sensory and language-related areas, redundant information is strongly significant in both hemispheres. Synergistic information in inferior frontal regions was also significant with more left-lateralized pattern. It should be noted that this pattern arises when perceived speech is natural unlike when task is challenging as in AV congruent condition (the main analysis), so it is highly likely that redundant information is stronger than synergistic information.

**S2 Fig.**
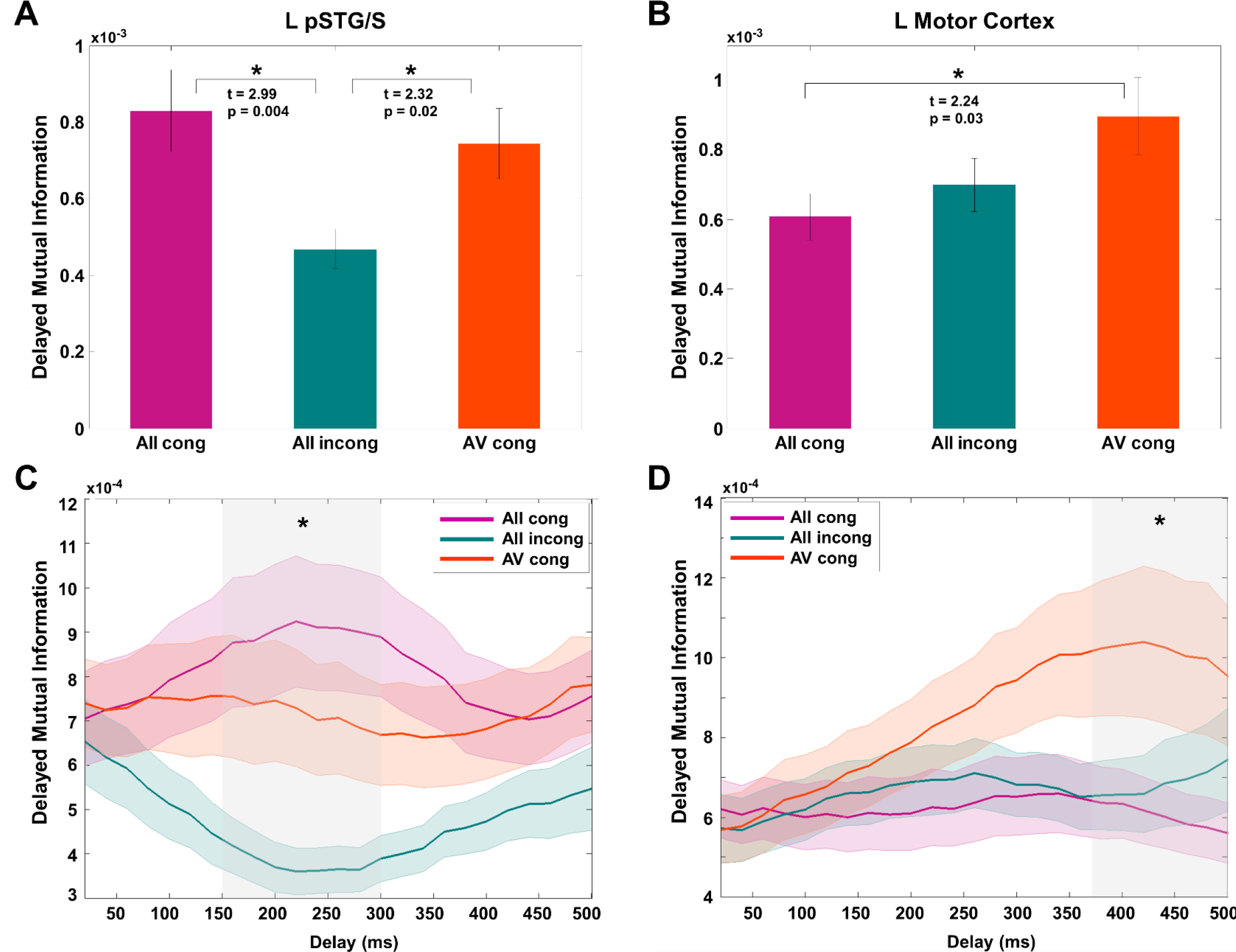
Left pSTG and left motor cortex differentially predict visual speech. A potential benefit of speech-entrained brain activity is the facilitation of temporal prediction of upcoming speech. We therefore investigated to what extent the different integration mechanisms in pSTG and motor cortex (reflected by differences in redundancy versus synergy) lead to differences in prediction. Since informativeness changed most strongly for the visual (speech) input signal (informative for ‘AV congruent’, less informative for ‘All congruent’), we expected strongest prediction effects for visual speech. We investigated prediction by means of delayed MI (see Methods for details) between theta phase in each brain area and later theta phase in visual speech. We tested delays from 0 ms to 500 ms in steps of 20 ms and then averaged the values across these delays. Interestingly, prediction of visual speech varied in both brain areas between conditions, but in different ways. Left pSTG predicts visual speech stronger in ‘All congruent’ and ‘AV congruent’ conditions than in incongruent condition (A; t = 2.99, *P* = 0.004 in ‘All congruent’ > ‘All incongruent’; t = 2.32, *P* = 0.02 in ‘AV congruent’ > ‘All incongruent’). Left motor cortex predicts visual speech stronger for ‘AV congruent’ than ‘All congruent’ (B; attention effect; t = 2.24, *P* = 0.03). When we unfolded these patterns in the temporal domain more interesting pattern emerged. The prediction mechanism in left pSTG operates in shorter temporal delays of 150-300 ms (C; *P* < 0.05), but left motor cortex is involved in longer temporal delays of 350 ms and above (D; *P* < 0.05). These findings suggest that left pSTG is mostly sensitive to congruent audiovisual speech (as demonstrated by redundancy in Fig 4) and best predicts visual speech when congruent audiovisual speech is available in the absence of distracting input. This happens fast at shorter delays with visual speech. However, this pattern is different for left motor cortex. Here, we see better prediction in the ‘AV congruent’ condition, when visual speech information is informative, attended and useful to resolve a challenging listening task. Thus it has rather slow temporal dynamics at delays greater than 350 ms. The prediction of auditory speech was not different between conditions. This is expected because the level of auditory attention is similar across conditions. Overall, this suggests that integration mechanisms in left pSTG are optimized for congruent audiovisual speech. This is consistent with results in Figs 4, 5 that show prominent redundancy in left pSTG. Left motor cortex instead seems to play an important role when greater attentional efforts are required and potential conflicts need to be resolved (as is the case for ‘AV congruent’ condition).

**S3 Fig.**
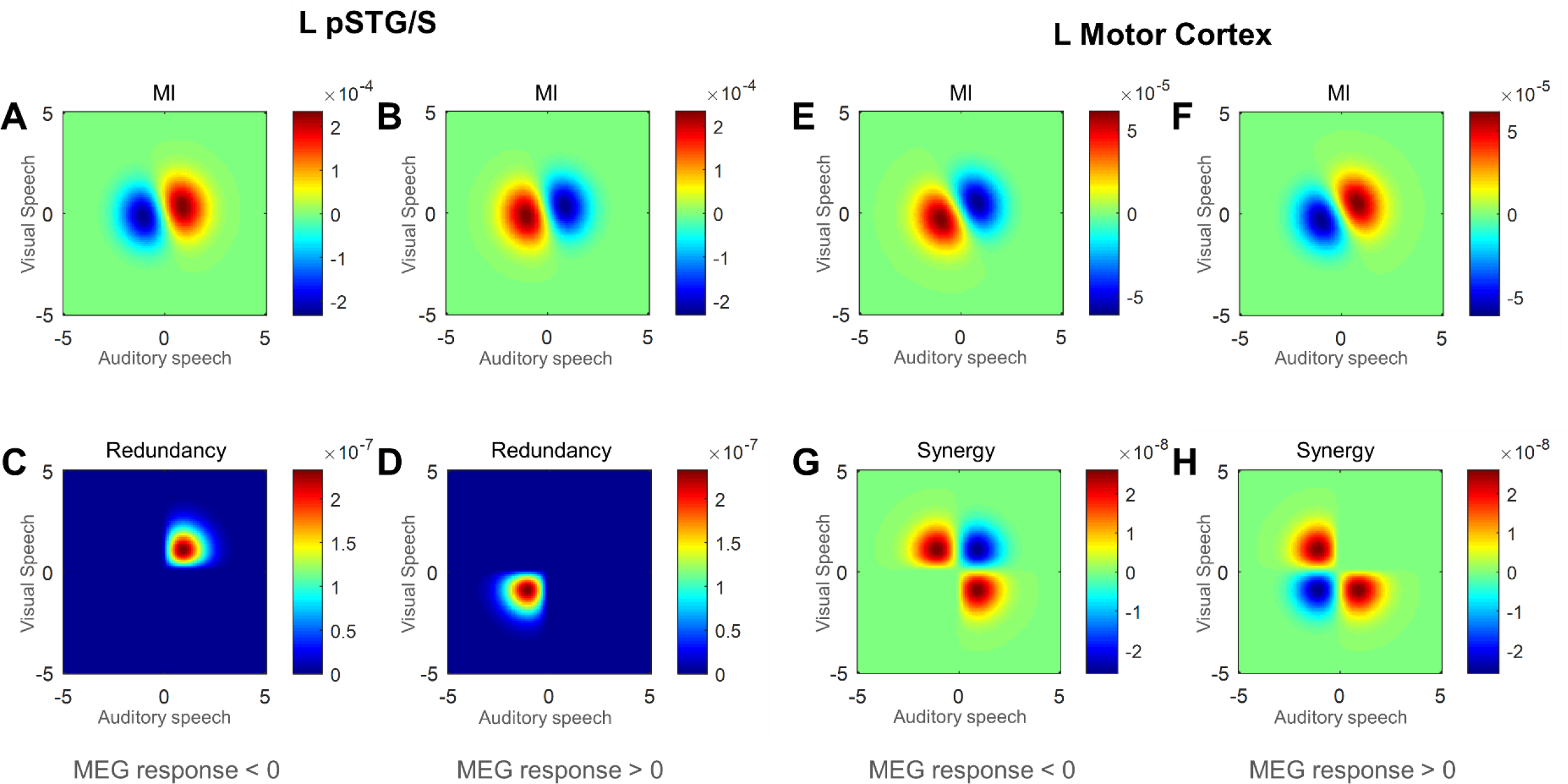
Local maps of redundancy and synergy in searching for mechanisms of AV speech interaction. This figure presents more detailed results from an analysis of interactions between auditory and visual speech signals as predictors of MEG signal in left superior temporal gyrus (pSTG) and left motor cortex. The selection of regions of interest was based on the results presented in Figures 3-5 that left pSTG features largely redundant interactions whereas left motor areas show predominantly synergistic interactions. As described in the Methods section, we estimate information quantities using Gaussian-Copula Mutual Information (GCMI) [30] which provides a robust semi-parametric lower bound estimator of mutual information, by combining the statistical theory of copulas with the closed form solution for the entropy of Gaussian variables. Crucially, this method performs well for higher dimensional responses as required for measuring three-way statistical interactions and allows estimation over circular variables like phase. Complex spectra from Hilbert-transformed signals of auditory speech, visual speech and each brain region were amplitude normalized, and the real and imaginary parts were rank-normalized. The covariance matrix describing Gaussian-Copula dependence was computed. Information and PID values are expectations over the space of joint values. To get more insight into the mechanisms underlying the information theoretic quantification, we can directly visualise the values which are summed in the expectation, often called local values [54]. While the main analysis involved 2D Hilbert transformed signals, for ease of visualisation we here consider just the 1D bandpass filtered signal. Joint MI can be written as:

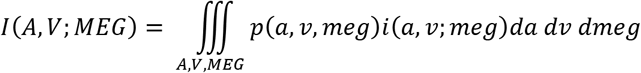

where 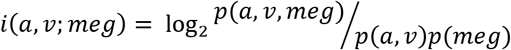 is the local information. We plot here the combined term *pi*, the local information quantity multipled by the probability of those values. This breaks down the overall quantity into the individual values which are integrated over the space. After copula normalization of a signal 0 corresponds to the median amplitude, so positive values on the x-and y-axis correspond to amplitude values above the median for auditory and visual speech signals, respectively. Here we study Mutual Information (MI) between auditory and visual speech for both regions of interest (upper row) and redundancy for left pSTG and synergy for left motor cortex (bottom row) averaged across all participants. In the ‘L pSTG’ plot, when MEG response < 0 (C), the redundancy comes from above median values of both auditory and visual speech. This shows that when both auditory and visual speech signals are high (above median), they redundantly suggest that MEG response is below the median value in the band. However, when both auditory and visual speech signals are below their median values, they redundantly suggest an above median MEG response (D). The ‘L Motor Cortex’ plot shows that when auditory and visual signals have opposite signs (i.e. above median value in one signal occurring with a below median value of the other signal, red diagonal nodes) (G,H) they synergistically inform about a co-occurring MEG response value. That is, knowing that A is high and V is low together (or know that V is high and A is low together), provides a better prediction of a specific MEG response value than would be expected if the evidence was combined independently. Here the negative node (blue) indicates a negative synergistic contribution to mutual information (sometimes called misinformation). Above/below median values can be interpreted as loud/quiet auditory speech and large/small lip movement, so low sound amplitude combined with small lip movement can produce larger response in the left pSTG whereas high sound amplitude combined with large lip movement can produce smaller response in the left pSTG. This seems highly plausible mechanism given the left pSTG’s role in AV integration that it has greater involvement when both speech are not physically strong enough. However, synergy shows a different pattern that regardless of high or low MEG response in the left motor cortex, the combination of low sound amplitude and large lip movement, or combination of high sound amplitude and small lip movement can produce synergistic information. This suggests synergistic information arises when the interaction comes out of unbalanced features of predictors that are complementary to each other.

**S4 Fig.**
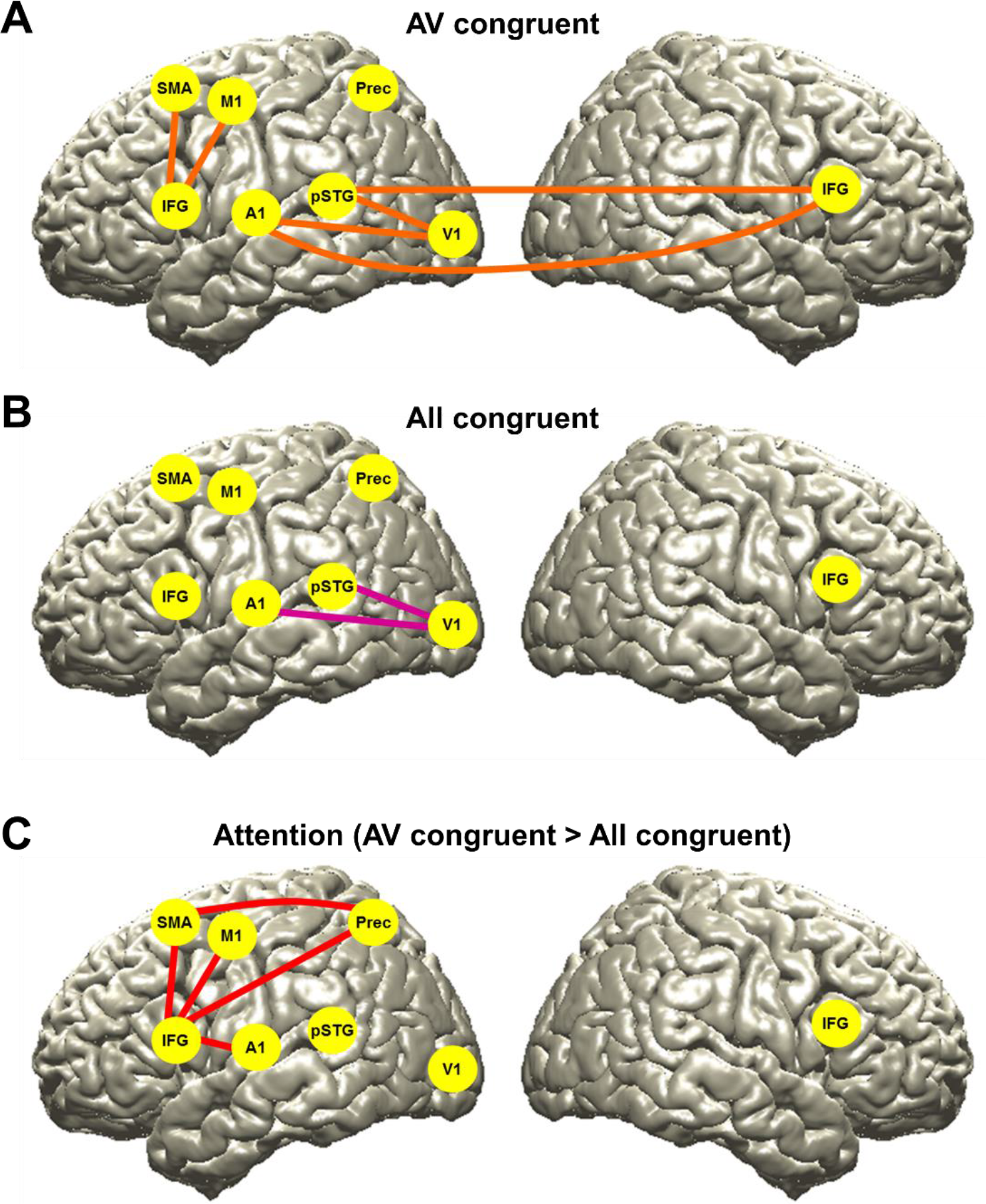
Synergy between brain regions predictive of visual speech. To further understand integration mechanism of audiovisual speech processing observed in redundant and synergistic interaction between multisensory speech signals predictive of brain activity, we computed PID differently in which redundant and synergistic interaction between brain regions predictive of speech signals (auditory or visual). We selected eight brain regions from the statistical contrasts for attention and congruence effects shown in Figs 4 and 5 (see Methods for details). The signals at the maximum coordinates were extracted from each region, PID was computed for each pair of brain regions predictive of auditory or visual speech signals. Each condition was compared to time-shifted surrogate data and between conditions. We found interesting results for synergistic interaction (but not for redundant information) between brain regions on visual speech for attention effect (‘AV congruent’ > ‘All congruent’). A. ‘AV congruent’ vs. surrogate data. Synergistic interaction between IFG (L)-M1, IFG (L)-SMA, A1-V1, A1-IFG (R), pSTG-IFG (R), pSTG-V1 were significant when predictive of visual speech. B. ‘All congruent’ vs. surrogate data. Synergistic interaction between A1-V1, pSTG-V1 were shown to be predictive of visual speech. C. ‘AV congruent’ vs. ‘All congruent’ (attention). When these two conditions were compared directly, synergistic interaction between IFG (L)-M1, IFG (L)-SMA, IFG (L)-A1, IFG (L)-Precuneus, SMA-Precuneus were observed to be predictive of visual speech (paired two-sided *t*-test; *P* < 0.05). However, we could not find any significant interaction (either redundancy or synergy) between these regions predictive of auditory speech. These results suggest that synergistic information interaction between the regions centering around left inferior frontal gyrus (BA44/BA45) and motor areas, which matches dorsal stream in speech processing [21], plays important role in attention to speech particularly visual speech when the task is challenging.

## References

1. Sumby WH, Pollack I. Visual Contribution to Speech Intelligibility in Noise. The Journal of the Acoustical Society of America. 1954;26(2).

2. McGurk H, MacDonald J. Hearing lips and seeing voices. Nature. 1976;264(5588):746–8. PubMed PMID: 1012311.

3. Grant KW, Seitz PF. The use of visible speech cues for improving auditory detection of spoken sentences. J Acoust Soc Am. 2000;108(3 Pt 1):1197–208. PubMed PMID: 11008820.

4. van Wassenhove V, Grant KW, Poeppel D. Visual speech speeds up the neural processing of auditory speech. Proc Natl Acad Sci U S A. 2005;102(4)1181–6. doi: 10.1073/pnas.0408949102. PubMed PMID: 15647358; PubMed Central PMCID: PMCPMC545853.

5. Park H, Kayser C, Thut G, Gross J. Lip movements entrain the observers’ low-frequency brain oscillations to facilitate speech intelligibility. Elife. 2016;5:e14521. doi: 10.7554/eLife.14521. PubMed PMID: 27146891; PubMed Central PMCID: PMCPMC4900800.

6. Zion Golumbic E, Cogan GB, Schroeder CE, Poeppel D. Visual input enhances selective speech envelope tracking in auditory cortex at a “cocktail party”. J Neurosci. 2013;33(4):1417–26. Epub 2013/01/25. doi: 10.1523/JNEUROSCI.3675-12.2013. PubMed PMID: 23345218; PubMed Central PMCID: PMC3711546.

7. Skipper JI, Devlin JT, Lametti DR. The hearing ear is always found close to the speaking tongue: Review of the role of the motor system in speech perception. Brain Lang. 2016;164:77–105. doi: 10.1016/j.bandl.2016.10.004. PubMed PMID: 27821280.

8. Calvert GA, Campbell R, Brammer MJ. Evidence from functional magnetic resonance imaging of crossmodal binding in the human heteromodal cortex. Curr Biol. 2000;10(11):649–57. PubMed PMID: 10837246.

9. Miller LM, D’Esposito M. Perceptual fusion and stimulus coincidence in the cross-modal integration of speech. J Neurosci. 2005;25(25):5884–93. doi: 10.1523/JNEUROSCI.0896-05.2005. PubMed PMID: 15976077.

10. Sekiyama K, Kanno I, Miura S, Sugita Y. Auditory-visual speech perception examined by fMRI and PET. Neurosci Res. 2003;47(3):277–87. PubMed PMID: 14568109.

11. Matchin W, Groulx K, Hickok G. Audiovisual speech integration does not rely on the motor system: evidence from articulatory suppression, the McGurk effect, and fMRI. J Cogn Neurosci. 2014;26(3):606–20. doi: 10.1162/jocn_a_00515. PubMed PMID: 24236768.

12. Beauchamp MS, Nath AR, Pasalar S. fMRI-Guided transcranial magnetic stimulation reveals that the superior temporal sulcus is a cortical locus of the McGurk effect. J Neurosci. 2010;30(7):2414–7. doi: 10.1523/JNEUROSCI.4865-09.2010. PubMed PMID: 20164324; PubMed Central PMCID: PMCPMC2844713.

13. Ding N, Melloni L, Zhang H, Tian X, Poeppel D. Cortical tracking of hierarchical linguistic structures in connected speech. Nat Neurosci. 2016;19(1):158–64. doi: 10.1038/nn.4186. PubMed PMID: 26642090; PubMed Central PMCID: PMCPMC4809195.

14. Giraud AL, Poeppel D. Cortical oscillations and speech processing: emerging computational principles and operations. Nat Neurosci. 2012;15(4):511–7. Epub 2012/03/20. doi: 10.1038/nn.3063. PubMed PMID: 22426255; PubMed Central PMCID: PMCPMC4461038.

15. Gross J, Hoogenboom N, Thut G, Schyns P, Panzeri S, Belin P, et al. Speech rhythms and multiplexed oscillatory sensory coding in the human brain. PLoS biology. 2013;11(12):e1001752. doi: 10.1371/journal.pbio.1001752. PubMed PMID: 24391472; PubMed Central PMCID: PMC3876971.

16. Ince RAA. Measuring multivariate redundant information with pointwise common change in surprisal. Entropy. 2017;19(7):318. Epub 29 June 2017. doi: 10.3390/e19070318.

17. Williams PL, Beer RD. Nonnegative Decomposition of Multivariate Information. arXiv:10042515v1. 2010.

18. Timme N, Alford W, Flecker B, Beggs JM. Synergy, redundancy, and multivariate information measures: an experimentalist’s perspective. Journal of computational neuroscience. 2014;36(2):119–40. doi: 10.1007/s10827-013-0458-4. PubMed PMID: 23820856.

19. Barrett AB. Exploration of synergistic and redundant information sharing in static and dynamical Gaussian systems. Phys Rev E Stat Nonlin Soft Matter Phys. 2015;91(5):052802. doi: 10.1103/PhysRevE.91.052802. PubMed PMID: 26066207.

20. Park H, Ince RA, Schyns PG, Thut G, Gross J. Frontal top-down signals increase coupling of auditory low-frequency oscillations to continuous speech in human listeners. Current biology: CB. 2015;25(12):1649–53. doi: 10.1016/j.cub.2015.04.049. PubMed PMID: 26028433; PubMed Central PMCID: PMCPMC4503802.

21. Hickok G, Poeppel D. The cortical organization of speech processing. Nature reviews Neuroscience. 2007;8(5):393–402. doi: 10.1038/nrn2113. PubMed PMID: 17431404.

22. Beauchamp MS. Statistical criteria in FMRI studies of multisensory integration. Neuroinformatics. 2005;3(2):93–113. doi: 10.1385/NI:3:2:093. PubMed PMID: 15988040; PubMed Central PMCID: PMCPMC2843559.

23. Campbell R, MacSweeney M, Surguladze S, Calvert G, McGuire P, Suckling J, et al. Cortical substrates for the perception of face actions: an fMRI study of the specificity of activation for seen speech and for meaningless lower-face acts (gurning). Brain Res Cogn Brain Res. 2001;12(2):233–43. PubMed PMID: 11587893.

24. Beauchamp MS, Lee KE, Argall BD, Martin A. Integration of auditory and visual information about objects in superior temporal sulcus. Neuron. 2004;41(5):809–23. PubMed PMID: 15003179.

25. Callan DE, Jones JA, Munhall K, Kroos C, Callan AM, Vatikiotis-Bateson E. Multisensory integration sites identified by perception of spatial wavelet filtered visual speech gesture information. J Cogn Neurosci. 2004;16(5):805–16. doi: 10.1162/089892904970771. PubMed PMID: 15200708.

26. Seltzer B, Pandya DN. Afferent cortical connections and architectonics of the superior temporal sulcus and surrounding cortex in the rhesus monkey. Brain research. 1978;149(1):1–24. Epub 1978/06/23. PubMed PMID: 418850.

27. Likert R. A technique for the measurement of attitudes. Archives of Psychology. 1932;22(140):1–55.

28. Brainard DH. The Psychophysics Toolbox. Spatial vision. 1997;10(4):433–6. PubMed PMID: 9176952.

29. Oostenveld R, Fries P, Maris E, Schoffelen JM. FieldTrip: Open source software for advanced analysis of MEG, EEG, and invasive electrophysiological data. Computational intelligence and neuroscience. 2011;2011:156869. Epub 2011/01/22. doi: 10.1155/2011/156869. PubMed PMID: 21253357; PubMed Central PMCID: PMC3021840.

30. Ince RA, Giordano BL, Kayser C, Rousselet GA, Gross J, Schyns PG. A statistical framework for neuroimaging data analysis based on mutual information estimated via a gaussian copula. Hum Brain Mapp. 2017;38(3):1541–73. doi: 10.1002/hbm.23471. PubMed PMID: 27860095.

31. Gross J, Baillet S, Barnes GR, Henson RN, Hillebrand A, Jensen O, et al. Good practice for conducting and reporting MEG research. NeuroImage. 2013;65:349–63. Epub 2012/10/11. doi: 10.1016/j.neuroimage.2012.10.001. PubMed PMID: 23046981.

32. Besl PJ, McKay ND. A method for registration of 3-D shapes. IEEE T Pattern Anal. 1992:239–56.

33. Nolte G. The magnetic lead field theorem in the quasi-static approximation and its use for magnetoencephalography forward calculation in realistic volume conductors. Physics in medicine and biology. 2003;48(22):3637–52. PubMed PMID: 14680264.

34. Gross J, Kujala J, Hamalainen M, Timmermann L, Schnitzler A, Salmelin R. Dynamic imaging of coherent sources: Studying neural interactions in the human brain. Proc Natl Acad Sci U S A. 2001;98(2)694–9. Epub 2001/02/24. doi: 10.1073/pnas.98.2.694. PubMed PMID: 11209067; PubMed Central PMCID: PMCPMC14650.

35. Chandrasekaran C, Trubanova A, Stillittano S, Caplier A, Ghazanfar AA. The natural statistics of audiovisual speech. PLoS computational biology. 2009;5(7):e1000436. doi: 10.1371/journal.pcbi.1000436. PubMed PMID: 19609344; PubMed Central PMCID: PMC2700967.

36. Smith ZM, Delgutte B, Oxenham AJ. Chimaeric sounds reveal dichotomies in auditory perception. Nature. 2002;416(6876):87–90. doi: 10.1038/416087a. PubMed PMID: 11882898; PubMed Central PMCID: PMC2268248.

37. Shannon CE. The mathematical theory of communication. Bell Syst Tech J. 1948;27:379.

38. Strong SP, de Ruyter van Steveninck RR, Bialek W, Koberle R. On the application of information theory to neural spike trains. Pac Symp Biocomput. 1998:621–32. PubMed PMID: 9697217.

39. Borst A, Theunissen FE. Information theory and neural coding. Nat Neurosci. 1999;2(11):947–57. doi: 10.1038/14731. PubMed PMID: 10526332.

40. Montemurro MA, Rasch MJ, Murayama Y, Logothetis NK, Panzeri S. Phase-of-firing coding of natural visual stimuli in primary visual cortex. Current biology: CB. 2008;18(5):375–80. doi: 10.1016/j.cub.2008.02.023. PubMed PMID: 18328702.

41. Rubino D, Robbins KA, Hatsopoulos NG. Propagating waves mediate information transfer in the motor cortex. Nat Neurosci. 2006;9(12):1549–57. doi: 10.1038/nn1802. PubMed PMID: 17115042.

42. Ince RA, Jaworska K, Gross J, Panzeri S, van Rijsbergen NJ, Rousselet GA, et al. The Deceptively Simple N170 Reflects Network Information Processing Mechanisms Involving Visual Feature Coding and Transfer Across Hemispheres. Cereb Cortex. 2016. doi: 10.1093/cercor/bhw196. PubMed PMID: 27550865; PubMed Central PMCID: PMCPMC5066825.

43. Schyns PG, Thut G, Gross J. Cracking the code of oscillatory activity. PLoS biology. 2011; 9(5)e1001064. doi: 10.1371/journal.pbio.1001064. PubMed PMID: 21610856; PubMed Central PMCID: PMCPMC3096604.

44. Kriegeskorte N, Mur M, Bandettini P. Representational similarity analysis - connecting the branches of systems neuroscience. Front Syst Neurosci. 2008;2:4. doi: 10.3389/neuro.06.004.2008. PubMed PMID: 19104670; PubMed Central PMCID: PMCPMC2605405.

45. King JR, Dehaene S. Characterizing the dynamics of mental representations: the temporal generalization method. Trends in cognitive sciences. 2014;18(4):203–10. doi: 10.1016/j.tics.2014.01.002. PubMed PMID: 24593982.

46. McGill WJ. Multivariate information transmission. Psychometrika. 1954;19(2):97–116. doi: 10.1007/BF02289159.

47. Wibral M, Priesemann V, Kay JW, Lizier JT, Phillips WA. Partial information decomposition as a unified approach to the specification of neural goal functions. Brain Cogn. 2017;112:25–38. doi: 10.1016/j.bandc.2015.09.004. PubMed PMID: 26475739.

48. Massey J. Causality, feedback and directed information. In: Proc Int Symp Information Theory Application (ISITA 1990). 1990:303–5.

49. Schreiber T. Measuring information transfer. Physical review letters. 2000;85(2):461–4. doi: 10.1103/PhysRevLett.85.461. PubMed PMID: 10991308.

50. Talairach J, Tournoux P. Co-Planar Stereotaxic Atlas of the Human Brain. NY: Thieme Medical Publishers; 1988.

51. Genovese CR, Lazar NA, Nichols T. Thresholding of statistical maps in functional neuroimaging using the false discovery rate. NeuroImage. 2002;15(4):870–8. doi: 10.1006/nimg.2001.1037. PubMed PMID: 11906227.

52. Eickhoff SB, Stephan KE, Mohlberg H, Grefkes C, Fink GR, Amunts K, et al. A new SPM toolbox for combining probabilistic cytoarchitectonic maps and functional imaging data. NeuroImage. 2005;25(4):1325–35. doi: 10.1016/j.neuroimage.2004.12.034. PubMed PMID: 15850749.

53. Tzourio-Mazoyer N, Landeau B, Papathanassiou D, Crivello F, Etard O, Delcroix N, et al. Automated anatomical labeling of activations in SPM using a macroscopic anatomical parcellation of the MNI MRI single-subject brain. NeuroImage. 2002;15(1):273–89. doi: 10.1006/nimg.2001.0978. PubMed PMID: 11771995.

54. Wibral M, Lizier JT, Priesemann V. Bits from Biology for Computational Intelligence. arXiv:14120291v1 [q-bioNC]. 2014.

